# Uncharted biosynthetic potential of the ocean microbiome

**DOI:** 10.1101/2021.03.24.436479

**Authors:** Lucas Paoli, Hans-Joachim Ruscheweyh, Clarissa C. Forneris, Satria Kautsar, Quentin Clayssen, Guillem Salazar, Alessio Milanese, Daniel Gehrig, Martin Larralde, Laura M. Carroll, Pablo Sánchez, Ahmed A. Zayed, Dylan R. Cronin, Silvia G. Acinas, Peer Bork, Chris Bowler, Tom O. Delmont, Matthew B. Sullivan, Patrick Wincker, Georg Zeller, Serina L. Robinson, Jörn Piel, Shinichi Sunagawa

**Author notes:** equally contributing second authors (Hans-Joachim Ruscheweyh, Clarissa C. Forneris).

## Abstract

Microbes are phylogenetically and metabolically diverse. Yet capturing this diversity, assigning functions to host organisms and exploring the biosynthetic potential in natural environments remains challenging. We reconstructed >25,000 draft genomes, including from >2,500 uncharacterized species, from globally-distributed ocean microbial communities, and combined them with ∼10,000 genomes from cultivated and single cells. Mining this resource revealed ∼40,000 putative biosynthetic gene clusters (BGCs), many from unknown phylogenetic groups. Among these, we discovered *Candidatus* Eudoremicrobiaceae as one of the most biosynthetically diverse microbes detected to date. Discrete transcriptional states structuring natural populations were associated with a potentially niche-partitioning role for BGC products. Together with the characterization of the first Eudoremicrobiaceae natural product, this study demonstrates how microbiomics enables prospecting for candidate bioactive compounds in underexplored microbes and environments.

## Introduction

Microbes drive global biogeochemical cycles, support food webs and underpin the health of animals and plants (Cavicchioli et al., 2019). Their metabolic and functional diversity represents a rich source of new drugs (Newman and Cragg, 2020) and revolutionary biotechnological applications (Knott and Doudna, 2018). Such discoveries have largely been facilitated by studying cultivable microbes under laboratory conditions. However, taxonomic surveys of natural environments have revealed that the vast majority of microbial life has not yet been cultivated (Lloyd et al., 2018). This cultivation bias has limited our ability to tap into much of the microbially-encoded functional diversity, despite its enormous potential for discovery of biosynthetic enzymes and bioactive compounds (Robinson et al., 2021).

Over the past decades, technological advances have enabled researchers to overcome these limitations by sequencing microbial DNA directly extracted from environmental samples (metagenomics). These efforts have provided a gene-level functional perspective to a previously taxon-centric exploration of microbiomes, *i*.*e*., microbial communities and their contained genetic material in a given environment (Qin et al., 2010; Sunagawa et al., 2015; Venter et al., 2004). Irrespective of the environment, metagenomic surveys have revealed that most of the functional diversity in different microbiomes had previously not been captured by reference genome sequences (REFs) from cultivated microbes (Nayfach et al., 2016). Placing this uncovered functional potential into a host genomic (*i*.*e*., genome-resolved) context has remained challenging, although doing so is critical to better understand organismal interactions, to study genomic variations, or to trac the source of metabolic activity (Sugimoto et al., 2019).

Recent advances in the field of microbiomics (*i*.*e*., the study of microbiomes) have enabled cultivation-independent, genome-resolved analyses of microbial communities. Algorithmic improvements have been used to group reconstructed genomic fragments from complex metagenomes into metagenome-assembled genomes (MAGs) (Albertsen et al., 2013). This approach has been instrumental in predicting, for example, the existence of subsequently isolated organisms that have provided additional clues on the origin of eukaryotic life (Imachi et al., 2020; Spang et al., 2015) and other yet uncharacterized soil lineages putatively encoding for new natural products (Crits-Christoph et al., 2018; Nayfach et al., 2020). Other technological developments, such as the isolation of single microbial cells followed by whole-genome amplification, have enabled the generation of single amplified genomes (SAGs) (Woyke et al., 2017). SAGs have, for example, provided insights into the limited clonality in natural populations of Key ocean bacteria (Kashtan et al., 2014) and led to the discovery of a metabolically rich group of sponge-associated bacteria, *Candidatus* Entotheonella, as a source of novel classes of candidate drugs (Wilson et al., 2014). More generally, large-scale efforts have vastly extended the phylogenomic representation of microbial diversity on Earth (Hug et al., 2016; Parks et al., 2018) and enabled genome-resolved explorations of various microbiomes (Almeida et al., 2020; Nayfach et al., 2020; Pachiadaki et al., 2019).

For the open ocean, however, genome-resolved data resources currently leave at least two-thirds of global metagenomic data unaccounted for, with even greater fractions in deeper waters and polar regions (Nayfach et al., 2020; Pachiadaki et al., 2019). Thus, the majority of the ocean microbiome along with its reservoir of biosynthetic potential for the production of uncharted bioactive natural products remains underexplored. Here, we integrated ocean microbial genomes from cultivation-dependent and independent methods to establish the most extensive phylogenomic and functional representation of the global ocean microbiome so far. We leveraged this resource to uncover a diverse array of biosynthetic gene clusters (BGCs), the majority of them from uncharacterized gene cluster families (GCFs). We further demonstrate the discovery potential of this community resource by identifying an uncharted family of bacteria that display the highest known diversity of BGCs in the pelagic oceans to date. By integrating environmental gene expression data, we shed light on the evolution and ecology of this newly identified group of biosynthetically ‘talented’ bacteria. Experimental validation of a novel ribosomally synthesized and post-translationally modified peptide (RiPP) demonstrates the viability of a microbiomics-driven strategy to access the ocean microbiome as a rich, yet largely untapped source of new natural products.

## Results and Discussion

### Phylogenomic representation of the ocean microbiome

To explore the biosynthetic potential of the ocean microbiome, we first sought to establish a global genome-resolved data resource focusing on its bacterial and archaeal constituents. To this end, we aggregated metagenomic data along with contextual information from 1,038 ocean water samples from 215 globally-distributed sampling sites (latitudinal range = 141.6°), several depth layers (from 1 to 5,600 meters deep, covering epi-, meso-, and bathypelagic zones), including two site-specific time series sampling programmes (Acinas et al., 2019; Biller et al., 2018; Salazar et al., 2019) (Figure 1A-B, Table S1). In addition to providing broad geographic coverage, these size-selectively filtered samples allowed for comparing different components of the ocean microbiome, including virus-enriched (<0.2 µm), prokaryote-enriched (0.2-3 µm), particle-enriched (0.8-20 µm) and virus-depleted (>0.2 µm) communities. We reconstructed a total of 26,293 predominantly bacterial and archaeal MAGs (Figure 1C, Figure S1). Specifically, we generated MAGs based on assemblies from individual, rather than pooled, metagenomic samples to prevent the collapsing of natural sequence variations across samples from different locations or different time points. In addition, we grouped genomic fragments based on their abundance correlation across large numbers of samples (between 58 and 610 samples, depending on the survey) (Methods). We found this step, which was omitted in previous large-scale MAGs reconstruction efforts (Nayfach et al., 2020; Pasolli et al., 2019), to significantly improve benchmar s of both the number (mean 2.7 times) and quality score (mean +20%) of reconstructed genomes (Figure S2) (Supplemental information). Overall, these efforts have increased the number of ocean water microbial MAGs by a factor of 4.5 (6 when counting high-quality MAGs only) compared to the most comprehensive MAG resource available to date (Nayfach et al. 2020; Methods). This set of newly created MAGs was combined with 830 manually-curated MAGs (Delmont et al., 2018), 5,969 SAGs (Klemetsen et al., 2018; Pachiadaki et al., 2019) and 1,707 REFs (Klemetsen et al., 2018) of marine bacteria and archaea into a combined collection of 34,799 genomes (Figure 1C).

**Figure 1:**
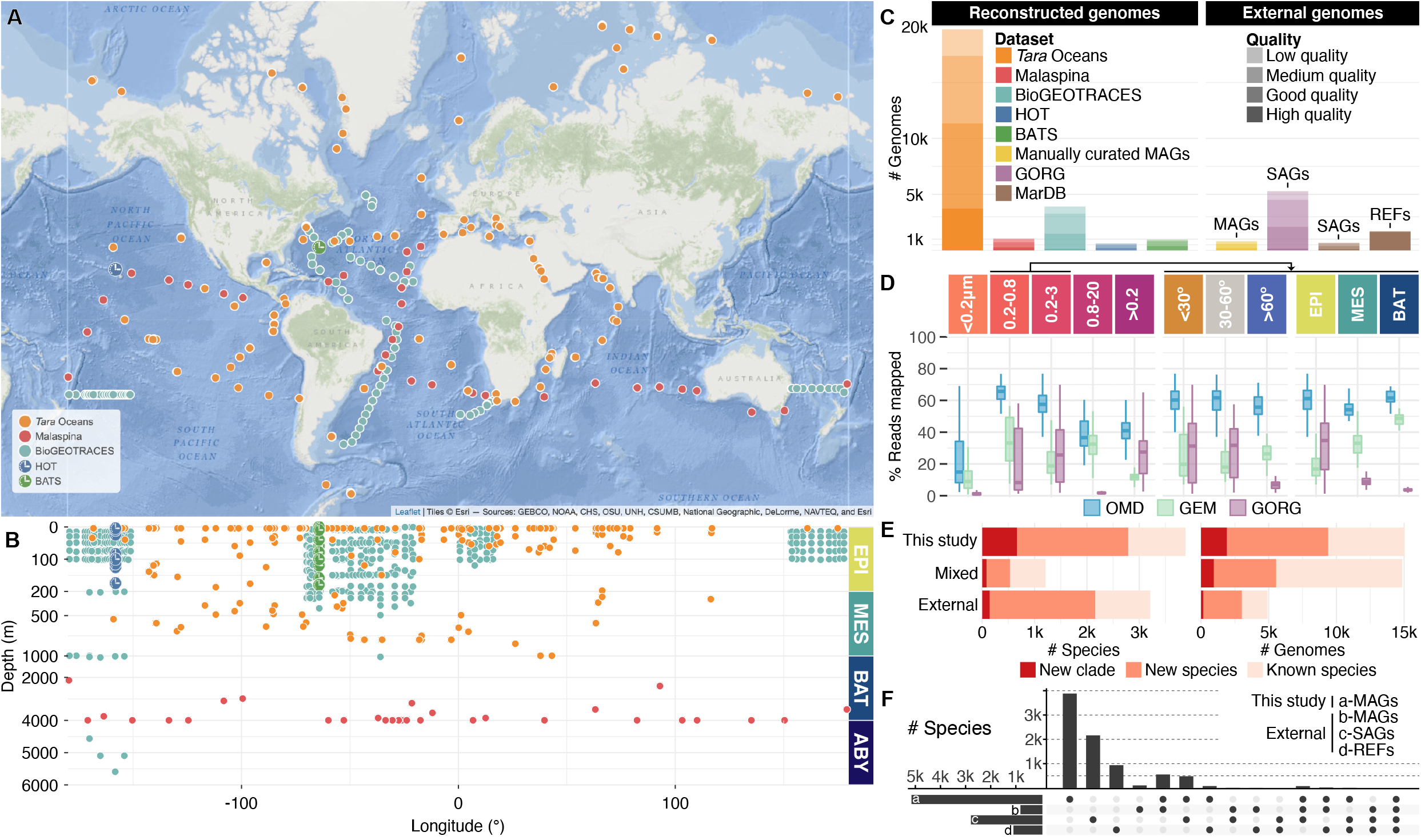
Reconstruction of MAGs at global scale fills ocean phylogenomic diversity gaps. (A) The aggregated set of metagenomes includes 1,038 ocean microbial community samples, which were collected at 215 globally-distributed sites (latitudinal and longitudinal range from 62°S to 79°N and 179°W to 179°E, respectively) in the context of different ocean expeditions and time series programmes. (B) The sample set covers a depth range from 1 to 5,601 meters representing major ocean depth layers. (C) Reconstructed MAGs varied in numbers and quality (Methods) across different data sets (color-coded). These MAGs were complemented with publicly available (external) genomes, including manually-curated MAGs (Delmont et al., 2018), single-amplified genomes (SAGs) (Klemetsen et al., 2018; Pachiadaki et al., 2019) and reference genomes from sequenced isolates (REFs) (Klemetsen et al., 2018) to compile the Ocean Microbiomics Database (OMD). (D) The OMD improves the genomic representation (mapping rates of metagenomic reads) of ocean microbial communities (from different size fractions) by a factor of two to three compared to previous reports solely based on SAGs (GORG) (Pachiadaki et al., 2019) or MAGs (GEM) (Nayfach et al., 2020), with a more consistent representation across depth and latitudes. (E) Grouping the OMD into species-level (95% average nucleotide identity) clusters identified a total of ∼8,300 species. Over half of these species were found to be uncharacterized, including a substantial fraction at the genus-level or above (new clades) based on taxonomic annotations using the Genome Taxonomy Database (GTDB) (release 89) (Parks et al., 2018). (D) Brea ing down the ∼8,300 species by genome type, we observed a high complementarity of MAGs, SAGs and REFs in capturing the phylogenomic diversity of the ocean microbiome. Specifically, 55%, 26% and 11% of the species were specific to MAGs, SAGs and REFs, respectively. HOT - Hawaiian Ocean Time Series; BATS - Bermuda Atlantic Time Series; EPI - epipelagic layer; MES - mesopelagic layer; BAT - bathypelagic layer; ABY - abyssopelagic layer; GEM - Genomes from Earth’s Microbiomes; GORG - Global Ocean Reference Genomes.

We next evaluated the newly established resource for its improved capacity to represent ocean microbial communities and to assess the impact of integrating different genome types. On average, we found it to capture about 40 to 60% of ocean metagenomic data (Figure 1D), corresponding to a two-to three-fold increase in coverage with a more consistent representation across depths and latitudes compared to previous reports solely based on MAGs (Nayfach et al., 2020) or SAGs (Pachiadaki et al., 2019). Furthermore, to obtain a systematic measure of the taxonomic diversity within the established collection, we annotated all genomes with the Genome Taxonomy Database (GTDB) Tool it (Methods) and clustered them using a 95% whole genome average nucleotide identity cutoff (Jain et al., 2018; Olm et al., 2020) to define 8,304 ‘species’-level clusters (species). Two thirds of these species (including new clades) were previously not represented in the GTDB and 2,790 of them were only uncovered by MAGs reconstructed in this study (Figure 1E). Additionally, we found the different genome types to be highly complementary, with 55%, 26% and 11% of the species being exclusively composed of MAGs, SAGs and REFs, respectively (Figure 1F). Furthermore, MAGs covered all 49 phyla detected in the water column, whereas SAGs and REFs represented only 18 and 11 of them, respectively. SAGs, however, better represented the diversity of the most abundant clades (Figure S3), such as the order Pelagibacterales (SAR11), with nearly 1,300 species covered by SAGs as opposed to only 390 by MAGs. Notably, REFs rarely overlapped with either MAGs or SAGs at the species level and represented >95% of the ∼1,000 genomes that were not detected in the set of open ocean metagenomes studied here (Methods), mostly owing to representatives that were isolated from other types of marine samples (*e*.*g*., sediments or host-associated). To enable a broad use by the scientific community, this ocean genomic resource, which also includes unbinned fragments (*e*.*g*., from predicted phages, genomic islands and fragments of genomes with insufficient data for MAG reconstruction) can be accessed alongside taxonomic and gene functional annotations as well as contextual environmental parameters at the Ocean Microbiomics Database (OMD): https://microbiomics.io/ocean/.

### Biosynthetic potential of the global ocean microbiome

Next, we set out to explore the richness and degree of novelty of the biosynthetic potential in the pelagic ocean microbiome. To this end, we first used antiSMASH to predict a total of 39,055 BGCs on all MAGs, SAGs and REFs detected in the set of 1,038 ocean metagenomes (Methods). Owing to the inherent redundancy of BGCs (*i*.*e*., the same BGC can be encoded in several genomes), we further clustered them into 6,907 non-redundant gene cluster families (GCFs) and 151 gene cluster clans (GCCs) (Table S2, Methods). To provide context, we compared these data with databases of computationally predicted (BiG-FAM) and experimentally validated (MIBIG 2.0) BGCs (Kautsar et al., 2020, 2021a).

We found the ocean microbiome to harbor a high diversity of RiPPs and other classes of candidate natural products at the GCC level (Figure 2A). Notably, aryl-polyenes, carotenoids, ectoines and siderophores, for instance, belonged to GCCs with wide phylogenomic distributions and high prevalence across ocean metagenomes, possibly indicative of widespread microbial adaptations to the ocean environment, including resistance to reactive oxygen species, oxidative- and osmotic-stress or uptake of iron (Supplemental information). This functional diversity contrasted with recent analyses of ∼1.2 M BGCs from ∼190,000 genomes deposited in the NCBI RefSeq database (BiG-FAM/RefSeq, hereafter RefSeq) (Kautsar et al., 2021a) that showed a dominance of non-ribosomal peptide synthetase (NRPS) and polyketide synthase (PKS) BGCs (Supplemental information). We additionally found that 44 GCCs were only remotely related to any RefSeq BGCs (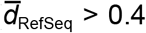, Figure 2A) (Methods), and 53 GCCs were only encoded in MAGs, highlighting the potential for discovery of novel chemistry within the OMD. Given that each of these GCCs is likely to represent highly diverse biosynthetic functions, we further analysed the data at the level of GCFs, which aim to provide a more fine-grained grouping of BGCs predicted to encode for similar natural products (Kautsar et al., 2021a). We found that 3,861 (56%) of the identified GCFs were not overlapping with RefSeq and that >97% of the GCFs were not represented in the most extensive database of experimentally validated BGCs (Kautsar et al., 2020) (Figure 2B). The majority of that novel diversity (78%) corresponded to predicted terpenes, RiPPs or other natural products, and a large fraction (47%) was encoded in phyla not generally known for their biosynthetic potential.

**Figure 2:**
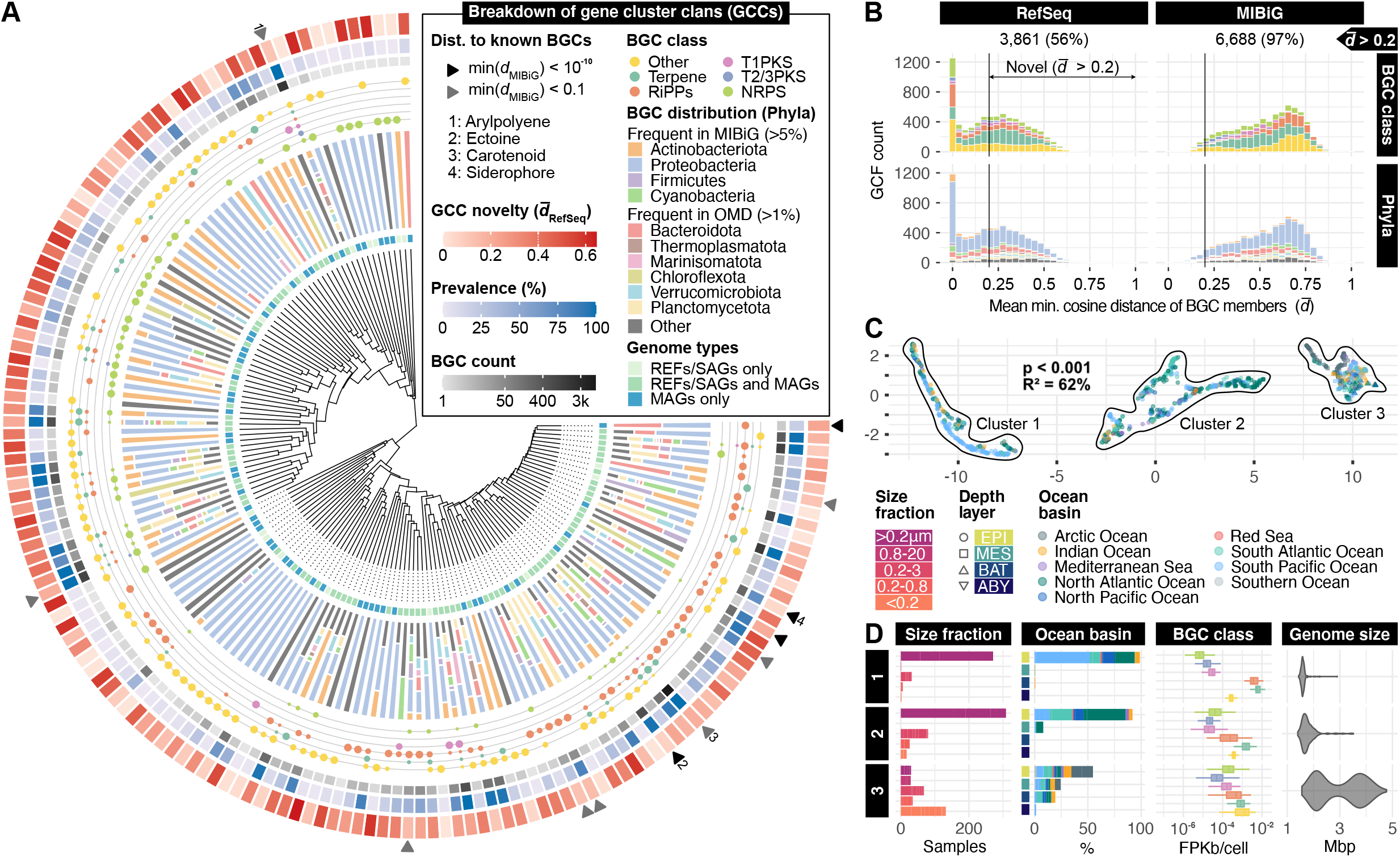
Novelty and structure of the ocean microbiome biosynthetic potential. The 39,055 predicted biosynthetic gene clusters (BGCs) were clustered into 6,907 non-redundant gene cluster families (GCFs) and 151 gene cluster clans (GCCs). (A) From the inner to outer layers: hierarchical clustering based on BGC distances of the 151 GCCs, 53 of which were only captured by MAGs (41 exclusive to this study). These GCCs comprise BGCs encoded by a variable number of phyla (shown as natural log-transformed frequencies) and belonging to different classes (shown as circles with sizes corresponding to frequency). Outer layers indicate the number of BGCs in a GCC, the prevalence of a GCC (as percentages of samples in which a GCC is detected) in the set of ocean metagenomes studied here (Methods) and the distance to BGCs in reference databases (mean min. cosine distance to RefSeq across the BGCs within a GCC). GCCs with BGCs closely related to characterized BGCs (from MIBiG) are highlighted by arrows. (B) The set of 6,907 GCFs was compared to computationally predicted (BiG-FAM/RefSeq) and experimentally validated (MIBIG 2.0) BGCs, uncovering 3,861 new 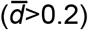 GCFs. Most (78%) of the new GCFs encode for RiPPs, terpenes and other putative natural products. (C) The abundances of GCFs (Methods) were used to compute distances between the 1,038 metagenomic samples. Using dimension reduction and density-based clustering (Methods), we identified three sample clusters. (D) These clusters were broken down by community origin, including size fractions, depth layers and ocean basins. We found significant differences (pairwise Wilcoxon tests) in BGC class abundances and average genome sizes (Methods) between the clusters (Table S2). RiPP - Ribosomally synthesized and Post-translationally modified Peptide; NRPS - Non-Ribosomal Peptide Synthetase; T1PKS - Type I Polyketide Synthase; T2/3PKS - Type II and III Polyletide Synthases.

We next sought to investigate the biogeographic structuring of the ocean biosynthetic potential by clustering samples based on the metagenomic abundance distribution of GCFs (Methods). Using dimensionality reduction and unsupervised density-based clustering (Methods), we identified three distinct clusters (PERMANOVA, p<0.001, Figure 2C). These clusters distinguished low latitude, epipelagic, pro aryote-enriched and virus-depleted communities, mostly from surface (cluster 1) or deeper sunlit waters (cluster 2), from polar, deep ocean, virus-enriched and particle-enriched communities (cluster 3) (Figure 2D). These differences are associated with higher abundances of RiPP and terpene BGCs in both clusters 1 and 2 as opposed to NRPS and PKS BGCs in cluster 1. We additionally found the clusters to contain samples with significantly different (Wilcoxon test, p<0.001) average genome sizes (Methods). Notably, streamlined genomes in clusters 1 and 2 were found to correlate with more compact BGCs (RiPPs, terpenes), while larger genomes in deeper and colder waters were also associated with larger BGCs (PKS, NRPS) (Figure S4, Figure 2D). Surprisingly, some of the largest average genome sizes, which are expected to positively correlate with cell size (Pachiadaki et al., 2019), were found in the virus-enriched (<0.22 µm) samples in cluster 3. Among the recovered MAGs, we found members of two new candidate orders within the Alpha- and Gammaproteobacteria, which were prevalent and often detected exclusively in this size fraction, with genome sizes ranging from 2 to 4.5 Mbp. Whether these MAGs represent other viable members of ultra-small bacteria (including pleomorphism or starvation forms) (Nakai, 2020) and/or content of gene transfer agents (Lang et al., 2012), rather than sampling artifacts remains to be elucidated. These observations complement previous gene-centric work (Lannes et al., 2019), which focused on ultra-small and genome-reduced microbes (Castelle et al., 2018) in a subset of virus-enriched *Tara* Oceans samples. Finally, we found well-studied tropical and epipelagic communities to appear as most promising sources of new terpenes, and the least explored communities (polar, deep, virus- and particle-enriched) to have the highest potential for the discovery of NRPS, PKS, RiPPs or other natural products (Figure S4).

### *Discovery of Candidatus* Eudoremicrobiaceae

We sought to map the phylogenomic distribution of the ocean microbiome biosynthetic potential, and to identify new BGC-rich clades. To this end, we placed the genomes in the standardized bacterial and archaeal phylogenomic trees of the GTDB (Parks et al., 2018), and overlayed the putative biosynthetic products they encode (Figure 3A). We found several BGC-rich clades (>15 BGCs) that are either well-known for their biosynthetic potential, such as Cyanobacteria (*Synechococcus*) and Proteobacteria (e.g., *Tistrella*) (Shah et al., 2017; Timmermans et al., 2017), or that have recently garnered attention for their natural products, such as Myxococcota (Sandaracinaceae), *Rhodococcus* and Planctomycetota (Ceniceros et al., 2017; Gregory et al., 2019; Wiegand et al., 2020), which were readily detected in ocean water samples (Methods). Interestingly, we found these clades to contain several unexplored lineages. For example, species with the richest biosynthetic potential within the Planctomycetota and Myxococcota phyla belonged to an uncharacterized candidate order and genus, respectively. Overall, this shows that the OMD provides access to previously uncharted phylogenomic information including for microbes that may represent novel targets for the discovery of enzymes and natural products.

**Figure 3:**
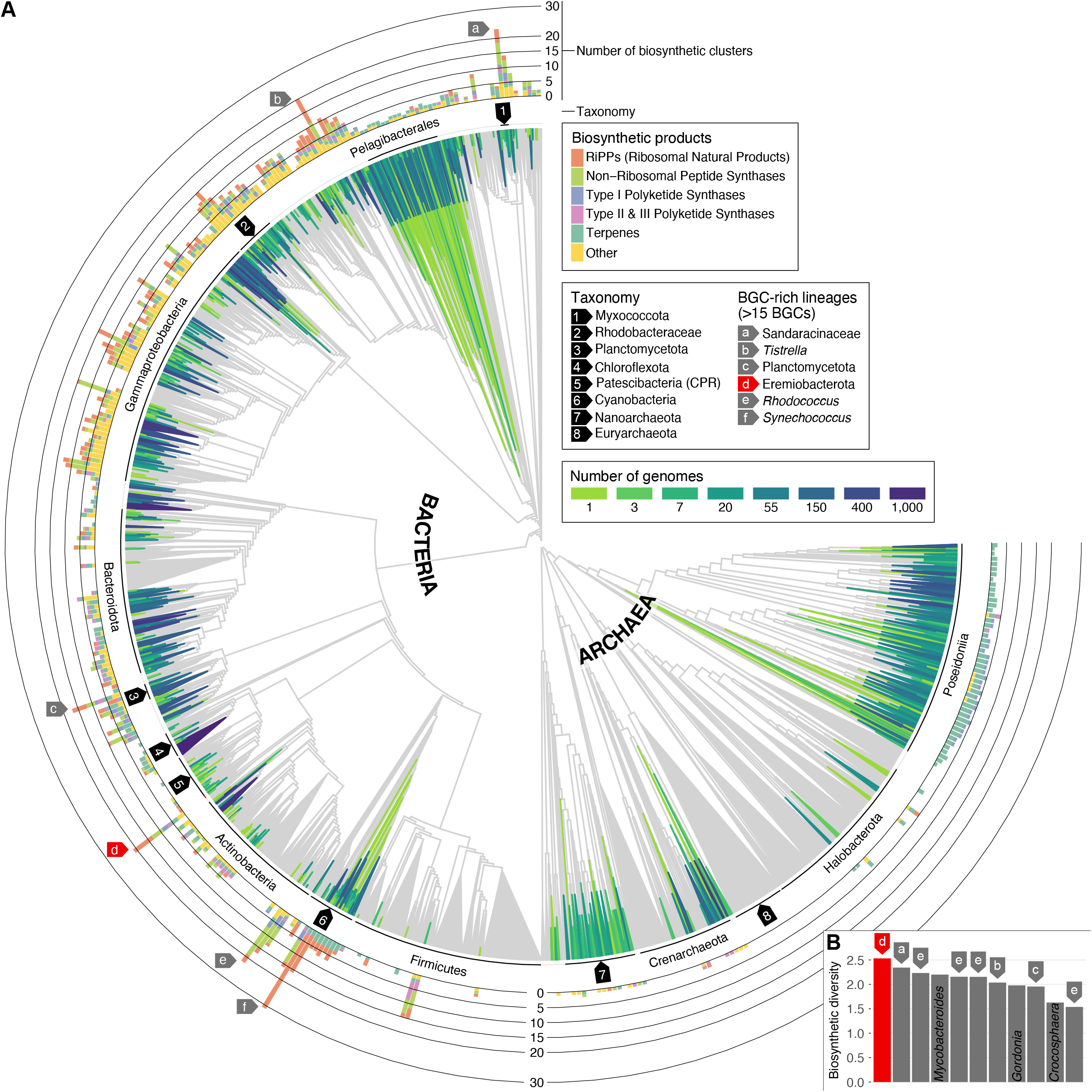
Phylogenomic distribution of the ocean microbiome biosynthetic potential. (A) Reconstructed MAGs, external MAGs, SAGs as well as REFs detected across the set of 1,038 ocean metagenomes were placed on the GTDB bac bone trees (Parks et al., 2018) to reveal the extent of phylogenomic coverage of the OMD. The taxonomy of some clades is indicated by arrows due to limited space. The number of BGCs corresponds to the highest number of predicted BGCs per genome in a given clade. Clades without any genome in the OMD are colored in gray. For visualization, the last 15% of the nodes are collapsed. Clades with >15 BGCs are denoted with arrows with the exception of *Mycobacteroides* and *Gordonia* (next to *Rhodococcus*) as well as Crocosphaera (next to *Synechococcus*). (B) Among the BGC-rich clades, we discovered an unknown species of *Ca*. Eremiobacterota to display the highest Shannon diversity index (y-axis) based on BGC-encoded predicted natural product types. Each bar represents the genome with the highest number of BGCs within a species. Species with a Shannon diversity index below 1.5 (incl. *Synechococcus* spp. and *Rhodococcus* spp.) are not shown.

To further characterize the BGC-rich clades, we set out to investigate not only the per genome total number of BGC-encoded natural products, but also their diversity (Figure 3B). We found the most biosynthetically diverse species to be exclusively represented by MAGs reconstructed in this study and to belong to the uncultivated phylum *Candidatus* Eremiobacterota. Notably, except for a few genomic studies (Ward et al., 2019; Woodcroft et al., 2018), this phylum has remained largely unexplored, and most strikingly, was so far unknown to contain any BGC-rich member. Eight MAGs with a nucleotide identity of >99% were reconstructed from deep (between 2,000 and 4,000 m) and particle-enriched (0.8-20 µm) ocean metagenomes collected by the Malaspina expedition (Acinas et al., 2019). Accordingly, we propose this species to be named ‘*Candidatus* Eudoremicrobium malaspinii’, after the nereid (sea nymph) of fine gifts in Greek mythology and the expedition.

*Ca*. E. malaspinii MAGs had no previously known relatives below the order level based on phylogenomic annotation (Parks et al., 2018), and belonged to the uncharacterized class UBP9 for which we now propose *Ca*. E. malaspinii as the type species and ‘*Candidatus* Eudoremicrobiia’ as its official name (Supplemental information). To search for closer relatives of this new species, we conducted targeted genome reconstructions in additional particle- and eukaryote-enriched metagenomic samples from the *Tara* Oceans expedition (Sunagawa et al., 2020). Briefly, we aligned metagenomic reads to *Ca*. E. malaspinii-related genomic fragments and assumed increased recruitment rates in a given sample to be indicative of the presence of additional relatives (Methods). As a result, we recovered 10 additional MAGs with the combined set representing five species across three genera within a newly defined family (*i*.*e*., *Ca*. Eudoremicrobiaceae). We then manually inspected and quality-controlled representative MAGs from each species (Supplemental information). We found these representatives to have larger genomes (>8 Mbp) with an actively transcribed (Figure S5) and richer biosynthetic potential (ranging from 14 to 22 BGCs per species) than in members of other *Ca*. Eremiobacterota clades (up to seven BGCs) from other environments (Figure 4A-C). The detection of this clade in less accessible (deep ocean) or eukaryote-enriched, rather than prokaryote-enriched samples, may explain why these bacteria and their unsuspected BGC diversity had remained elusive in the context of natural products research.

**Figure 4:**
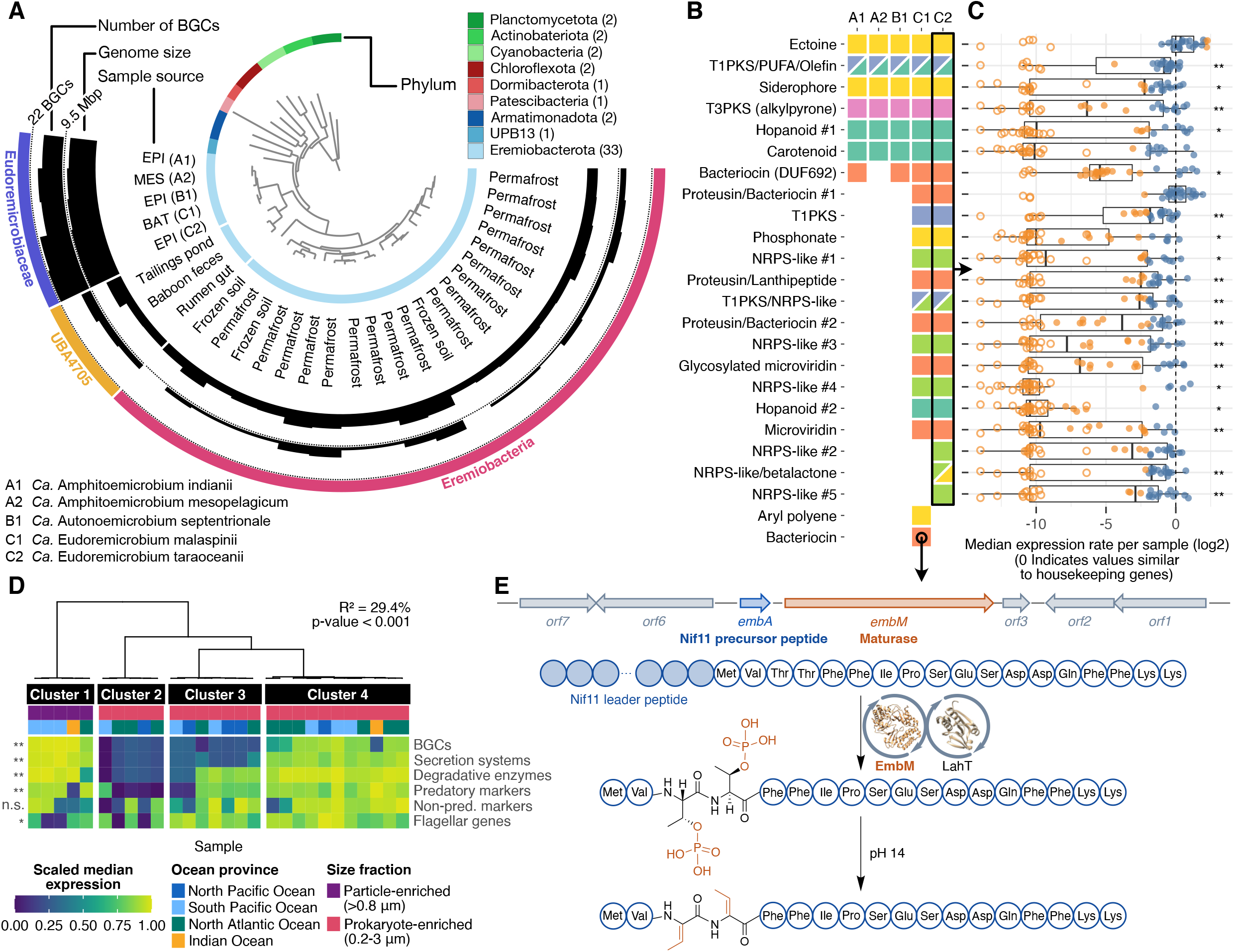
Discovery of an unsuspected, BGC-rich family in the phylum Ca. Eremiobacterota. (A) Phylogenomic placement of the five *Ca*. Eudoremicrobioceae spp. revealed that large genome sizes and high BGC-richness are specific to the discovered ocean lineage. The phylogenomic tree includes all *Ca*. Eremiobacterota MAGs available at GTDB (release 89) and representatives from additional phyla (number of genomes indicated in parentheses) for evolutionary context (Methods). The outermost layer indicates family- (*Ca*. Eudoremicrobiaceae and UBA4705) and class-level (*Ca*. Eremiobacteria) taxonomy. The five species described in this study are denoted by an alphanumeric code and a proposed binomial name (Supplemental information). (B) *Ca*. Eudoremicrobiaceae spp. share a core of seven BGCs. The missing BGC from clade A2 is most probably due to the incompleteness of the representative and only MAG available for this species (Table S3). This core potential is likely associated with osmotic stress (ectoine), iron uptake (siderophore) and membrane fluidity (PUFA, al ylpyrone) (Supplemental information). By contrast, *Ca*. Eudoremicrobium spp. (Clade C) collectively contain 17 specific BGCs including 10 NRPS and PKS as well as 7 RiPPs. BGCs specific to *Ca*. Amphithoemicrobium spp. and *Ca*. Autonoemicrobium spp. (Clade A and B respectively) are not displayed. Product classes are colored according to Figure 3. (C) All BGCs encoded by *Ca*. E. taraoceanii were found to be expressed across 623 metatranscriptomes sampled by *Tara* Oceans. Filled circles indicate active transcription. Orange data points indicate values below or above a log2-fold change from the expression rate of house eeping genes (Methods). (D) Four discrete expression states explained 29.4% of the overall transcriptomic variance (p<0.001, PERMANOVA) across *Ca*. E. taraoceanii populations. One state (cluster 1) was exclusive to larger organismal size fractions. Leafs represent transcriptomic profiles and the dendrogram represents dimensionality-reduced distances (Methods). Genes associated with BGCs, secretion systems, degradative enzymes and predatory mar ers were differentially expressed across the states and represented the most discriminatory categories compared to 200 KEGG pathways (Table S4). (E) *In vitro* heterologous expression of a bacteriocin biosynthetic cluster specific to the deep ocean species *Ca*. E. malaspinii led to the production of a di-phosphorylated product, which upon chemical elimination resulted in a new linear dehydrated product. FDR-corrected p-values (Kruskal-Wallis test) (*p<0.05, **p<0.01) correspond to differentially expressed BGCs (panel C) or functional groups (panel D) across the four clusters shown in panel D.

### *Putative ecology of Ca*. Eudoremicrobium spp

To better understand the ecology of *Ca*. Eudoremicrobiaceae spp. in the global oceans, we sought to explore computational trait and lifestyle predictions (Methods) along with biogeographic patterns of their abundance and gene expression. The results suggest *Ca*. Eudoremicrobiaceae representatives to be putatively gram-negative, motile, heterotrophic bacteria. Members of this family harbor a rich repertoire of degradative enzymes as well as diverse secretion systems, which, along with their biosynthetic potential, could represent the genomic evidence for predatory behavior (Perez et al., 2016). We found support for this hypothesis in high predatory index (Pasternak et al., 2013) values for *Ca*. Eudoremicrobium spp., which scored even higher than the *Bdellovibrio* spp. that serve as models for predatory specialization in bacteria (Table S3). *Ca*. Eudoremicrobiaceae accounted locally for up to 6% of the microbial community and were found to be prevalent in most oceanic basins as well as throughout the water column (Figure S6), making them a numerically important component of the global ocean microbiome.

To gain additional information on the putative lifestyle of these bacteria, we examined the biogeographic patterns of gene expression across the *Tara* Oceans metatranscriptomic samples (Sunagawa et al., 2020) for ‘*Candidatus* Eudoremicrobium taraoceanii’, which was found to be the most prevalent *Ca*. Eudoremicrobiaceae species in the surface oceans (Figure S6) and to display little gene content variation (Figure S7). Through dimensionality reduction and unsupervised density-based clustering, we determined that 29.4% of the transcriptome variance could be explained by four discrete clusters (p-value <0.001) reflecting distinct transcriptional states (Figure 4D). One of these states corresponds to the gene expression profile of *Ca*. Eudoremicrobium taraoceanii in all particle- and eukaryote-enriched samples (all samples >0.8 µm: i.e., 0.8-5, >0.8 and 5-20 µm). The remaining transcriptional states were all derived from prokaryote-enriched samples (0.2-3 µm). Across all four states, there was no clear separation by geographic origin. Combined with the predicted traits (motile, heterotrophic, predatory), these findings suggest that *Ca*. Eudoremicrobium spp. may alternate between a free-living (prokaryote-enriched) and a particulate organic matter-associated state.

We further sought to test whether these states may be lin ed to a secondary metabolite-driven predatory behavior as reported for some antibiotics-producing bacteria (Xiao et al., 2011). To explore this hypothesis, we identified differentially expressed functional groups across the four states (Supplemental information) and found BGCs, secretion systems, degradation enzymes and predatory markers to be among the most discriminative ones (Table S4). Specifically, the expression states of these groups were on average highest in the particle-enriched size fractions, although we also found them to be expressed in one of the free-living states (Figure 4D). By contrast, flagellar genes were more highly expressed in prokaryote-enriched fractions, further supporting the idea of particle-attached vs free-living lifestyles of these bacteria. Together, these observations suggest a strong association between the ecology of *Ca*. E. taraoceanii and the expression of its BGCs, which may imply a niche-partitioning role for the candidate natural products they encode.

### *The uncharted metabolic potential of Ca*. Eudoremicrobiaceae

To further explore the remarkable biosynthetic potential within *Ca*. Eudoremicrobiaceae, we contrasted its core and clade-specific BGCs. We found the five species to share seven clusters encoding putative natural products most likely involved in the regulation of osmotic stress, iron uptake and membrane fluidity (Figure 4B) (Supplemental information). Nonetheless, the *Ca*. Eudoremicrobium genus stood out as its members were found to encode for up to 15 additional BGCs. Among them, 10 were identified as NRPS and PKS clusters, including a particularly large (over 70 bp) and architecturally complex type I PKS/NRPS hybrid. An unusual feature of this cluster is the presence of terminal reductase domains in both of the two NRPS proteins. Notably, the majority of characterized natural products modified by these domains have shown varied and enhanced bioactivity such as protease inhibition and antitumoral properties (Mullowney et al., 2018).

In addition, we found *Ca*. Eudoremicrobium MAGs to contain up to seven RiPP BGCs spanning multiple peptide classes (Montalban-Lopez et al., 2020), including three proteusin BGCs (more than in any organism analysed so far). Proteusins are of particular biotechnological interest owing to their varied bioactivities and the expected density and diversity of chemical modifications installed by enzymes encoded in relatively short BGCs (Freeman et al., 2012). One such *Ca*. Eudoremicrobium BGC is of exceptional complexity as it encodes over 10 maturation enzymes, including a lanthionine synthetase and three epimerases that likely modify a precursor peptide along with the complete genetic machinery for cleavage and transport of the mature proteusin. The predicted presence of o-amino acids and (methyl)lanthionine rings resulting from the first half of the cluster suggests some shared structural features (Supplemental information) to the recently characterized antiviral landornamides (Bosch et al., 2020). The homology of some of *Ca*. Eudoremicrobium BGCs to clusters producing known cytotoxic natural products (Supplemental information), along with their reported enriched expression in particle- and eukaryote-enriched fractions and predicted predatory behavior could suggest eukaryotic preys, as reported for specialized metabolites-producing predators such as *Myxococcus* (Perez et al., 2016).

Due to its ecological and enzymatic particularity (Figure 4B), we selected a predicted novel bacteriocin BGC 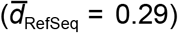 of *Ca*. E. malaspinii for experimental characterization. This BGC encodes a Nif11-type precursor peptide, EmbA, and a protein containing a predicted dehydration domain homologous to LanM-type lanthionine synthetases, EmbM, as a sole identified post-translationally modifying enzyme (Figure 4E). Heterologous expression of this cluster in *Escherichia coli* and *Microvirgula aerodenitrificans* and *in vitro* enzyme activity assays (Methods) resulted in high conversion rates (80%) to modified peptides. Curiously, the products were efficiently phosphorylated at both threonines, confirming the predicted enzyme activity, but the expected second half of the reaction resulting in formation of dehydrated threonines was not observed, although we could obtain the dehydrated hypothetical end-products by chemical elimination (Supplemental information). Both products were inactive against several tested pathogenic and marine bacterial strains as well as fungal strains (Methods). This was not entirely unexpected considering that bacteriocins often exhibit activity only against strains closely related to the producer (Soltani et al., 2021) and that *Ca*. Eremiobacterota and most of their closely related phyla have no cultivated representatives. Nevertheless, these synthetic biology efforts have enabled us to begin the functional characterization of the richness, diversity and unusual architectures identified in *Ca*. Eudoremicrobium BGCs.

## Conclusions

The present work has unveiled the extent of microbially-encoded biosynthetic potential and its genomic context in the global ocean microbiome, and facilitates future research by ma ing the generated resources available to the scientific community (https://microbiomics.io/ocean/). Finding the majority of both its phylogenomic and functional novelty to be accessible only through the reconstruction of MAGs, particularly in less explored microbial communities, could direct future bioprospecting efforts. Although the focus here has been on *Ca*. Eudoremicrobiaceae as a particularly biosynthetically ‘talented’ lineage, many of the predicted BGCs within other underexplored microbial groups are likely to produce compounds with novel activities. These activities may be ey to better understand microbial interactions and population structures in the natural environment, and thus motivate studying the diversity and function of BGCs, not only for their biotechnological relevance, but also within an eco-evolutionary context.

## Acknowledgements

We are greatly indebted to the work of *Tara* Oceans, which has been driven by a collaborative engagement of the Tara Ocean Foundation, countless scientific project members and associated institutional partners. We are grateful for the availability of the data generated in the context of the BioGEOTRACES, Malaspina, HOT and BATS projects, which formed the basis of the present work. We further thank the ETH Zurich HPC facilities for computational support. This article is the contribution number TBD of *Tara* Oceans. This work was supported by the ETH and the Helmut Horten Foundation and by funding from the Swiss National Foundation through project grants 205321_184955 to S.S., 205320_185077 to J.P., the NCCR Microbiomes (51NF40_180575), and the Peter and Traudl Engelhorn Foundation (to C.C.F.). S.L.R. was supported by an ETH Zurich Postdoctoral Fellowship 20-1 FEL-07. M.L, L.M.C. and G.Z. were supported by EMBL Core Funding and the German Research Foundation (DFG, Deutsche Forschungsgemeinschaft, project no. 395357507 - SFB 1371 to G.Z.). M.B.S. was supported by the NSF grant OCE#1829831. C.B. was supported by the European Research Council (ERC) under the European Union’s Horizon 2020 research and innovation programme (grant agreement Diatomic, No. 835067). S.G.A. was supported by the Spanish Ministry of Economy and Competitiveness (CTM2017-87736-R).

## Author contributions

Conceptualization: L.P., J.P., S.S. - Methodology, Validation and Data Curation: L.P., H.J.R., C.C.F., S.K., S.L.R., J.P., S.S. - Software: L.P., H.J.R., S.K., Q.C. - Investigation and Formal Analysis: L.P., H.J.R., C.C.F., S.K., Q.C., G.S., A.M., D.G., M.L., L.M.C., P.S., T.O.D., S.L.R., J.P., S.S. - Visualization: L.P., C.C.F., S.K., S.L.R. - Writing - Original Draft: L.P., S.S. - Writing - Review & Editing: H.J.R., C.C.F., S.K., A.A.Z., D.R.C., S.G.A., P.B., C.B., T.O.D., M.B.S., P.W., G.Z., S.L.R., J.P. - Funding Acquisition, Resources, Project Administration and Supervision: J.P., S.S..

