## Supplementary material for "Uncharted biosynthetic potential of the ocean microbiome": Methods and Supplemental Information

#### RESOURCE AVAILABILITY

*Lead contact*

*Materials Availability*

Material generated in this study is available upon request.

*Data and Code Availability*

Metagenomic assemblies are available at the ENA (TBD), additional supporting data have been deposited on Zenodo (https://doi.org/10.5281/zenodo.4474310), and code used for the analyses performed in this study is accessible at https://microbiomics.io/ocean/.

#### METHODS DETAILS

*Metagenomic data selection, assembly and binning*

Metagenomic datasets from major oceanographic surveys and time-series studies were included to maximize the coverage of global ocean microbial communities across ocean basins, depth layers and time. These datasets (Table S1, Figure 1) included metagenomes from samples collected by: *Tara* Oceans (virus-enriched, n=190, and prokaryote-enriched, n=180) (Salazar et al., 2019; Sunagawa et al., 2015), BioGEOTRACES expeditions (n=480), the Hawaiian Ocean Time-series (HOT, n=68), the Bermuda-Atlantic Time-series Study (BATS, n=62) (Biller et al., 2018) and the Malaspina expedition (n=58) (Acinas et al., 2019). Sequencing reads from all metagenomes were quality filtered using BBMap (v.38.71) by removing sequencing adapters from the reads, removing reads that mapped to quality control sequences (PhiX genome) and discarding low quality reads using the parameters *trimq=14, maq=20, maxns=0* and *minlength=45*. Downstream analyses were performed on quality-controlled reads, or, if specified, merged quality-controlled reads (bbmerge.sh *minoverlap=16*). Quality-controlled reads were normalized (bbnorm.sh *target=40, mindepth=0*) before they were assembled with metaSPAdes (v3.11.1 or v3.12 if required) (Nurk et al., 2017). The resulting scaffolded contigs (hereafter scaffolds) were finally filtered by length (≥1 kbp).

The 1,038 metagenomic samples were grouped into several sets, and for each sample set, the quality-controlled metagenomic reads from all samples were individually mapped against the scaffolds of each sample, resulting in the following numbers of pairwise readset to scaffold mappings: *Tara* Oceans virus-enriched (190*190), prokaryote-enriched (180*180),

BioGEOTRACES, HOT and BATS (610*610) and Malaspina (58*58). Mapping was performed using the Burrows-Wheeler-Aligner (BWA) (v0.7.17-r1188) (Li and Durbin, 2009), allowing the reads to map at secondary sites (with the *-a* flag). Alignments were filtered to be at least 45 bases in length, with an identity of ≥97% and covering ≥80% of the read sequence. The resulting BAM files were processed using the jgi_summarize_bam_contig_depths script of MetaBAT2 (v2.12.1) (Kang et al., 2019) to provide within- and between-sample coverages for each scaffold. The scaffolds were finally binned by running MetaBAT2 on all samples individually with parameters *--minContig 2000* and *--maxEdges 500* for increased sensitivity. To test for the effect of abundance correlation, a randomly selected subsample of the metagenomes (10 from each of the two *Tara* Ocean datasets, 10 from BioGEOTRACES, five from each time-series and five from Malaspina) was additionally binned only using within-sample coverage information (Supplemental information).

*Selection of additional genomes*

Additional (external) genomes were included in downstream analyses, namely 830 manually-curated MAGs from a subset of the *Tara* Oceans dataset (Delmont et al., 2018), 5,287 Single-cell Amplified Genomes (SAGs) from the GORG dataset (Pachiadaki et al., 2019) as well as 1,707 isolate reference genomes (REFs) and 682 SAGs from the MAR databases (MarDB v4) (Klemetsen et al., 2018). For the MarDB dataset, genomes were selected based on the available metadata if the sample type matched the following regular expression: ‘[S|s]ingle.?[C|c]ell|[C|c]ulture|[I|i]solate’.

*Quality evaluation of metagenomic bins and external genomes*

The quality of each metagenomic bin and external genome was evaluated using both the ‘lineage workflow’ of CheckM (v1.0.13) and Anvi'o (v5.5.0) (Eren et al., 2015; Parks et al., 2015). Metagenomic bins and external genomes were kept for downstream analyses if either CheckM/Anvi'o reported a completeness/completion ≥50% and a contamination/redundancy ≤10%. This notably allowed for the identification of 148 Eukaryotic MAGs by Anvi'o. These metrics were then aggregated into a mean completeness (mcpl) and a mean contamination (mctn) value to classify genome quality according to community standards (Bowers et al., 2017) as follows: high quality: mcpl≥90% and mctn≤5%; good quality: mcpl≥70% and mctn≤10%; medium quality: mcpl≥50% and mctn≤10%; low quality: mcpl≤90% or mctn≥10%. Filtered genomes were further attributed quality scores (Q and Q’) as follows: Q = mcpl - 5*mctn; and Q’ = mcpl - 5*mctn + mctn*(Strain Heterogeneity)/100 + 0.5*log(N50) (as implemented in dRep (Olm et al., 2017)).

*Species-level clustering of the genome collection and comparison to other resources*

To allow for comparative analyses between different data resources and genome types (MAGs, SAGs and REFs), the full set of 34,799 genomes was dereplicated based both on whole genome average nucleotide identity (ANI) using dRep (v2.5.4) (Olm et al., 2017) with a 95% ANI threshold (Jain et al., 2018; Olm et al., 2020) (*-comp 0 -con 1000 -sa 0.95 -nc*

*0.2*) and single-copy marker genes using SpecI (Mende et al., 2013), providing species-level clustering of the genomes. A representative genome was selected based on the maximum quality score defined above (Q') for each of the dRep clusters, which were considered as a proxy for species membership.

To estimate mapping rates, all 1,038 metagenomic readsets were mapped against the 34,799 genomes included in the Ocean Microbiomics Database (OMD) using BWA (v0.7.17-r1188, *-a*). Quality-controlled reads were mapped in single-end mode and the resulting alignments were filtered to keep only alignments ≥45 bp in length and an identity ≥95%. The per-sample mapping rate is the percentage of reads that remained after filtering divided by the total number of quality-controlled reads. Using the same method, each of the 1,038 metagenomes was downsampled to 5 M inserts and mapped to the GORG SAGs within the OMD and all of the GEM (Nayfach et al., 2020). The number of MAGs in the GEM catalog (Nayfach et al., 2020) that were recovered from ocean waters was determined based on a keyword query on the source of metagenomes, selecting for ocean water samples (as opposed to marine sediments, for example). Specifically, we selected 'Aquatic' as the 'ecosystem_category', 'Marine' as the 'ecosystem_type' and filtered 'habitat' for 'Deep ocean', 'Marine', 'Marine Oceanic', 'Pelagic marine', 'Seawater', 'marine', 'seawater', 'surface seawater', 'Surface seawater'. This resulted in 5,903 MAGs (734 high quality), distributed across 1,823 OTUs (here species).

*Taxonomic and functional genome annotation*

Prokaryotic genomes were taxonomically annotated using GTDB-Tk (v1.0.2) (Chaumeil et al., 2019) with default parameters against the GTDB r89 release (Parks et al., 2018). Anvi'o was used to identify eukaryotic genomes based on domain prediction and completion ≥50% and redundancy ≤10%. The taxonomic annotation of a species is defined as the one of its representative genome. Excluding eukaryotes, each genome was functionally annotated by first calling complete genes using prokka (v1.14.5) (Seemann, 2014) with the '*Archaea'* or '*Bacteria'* parameter specified as appropriate, which also reported non coding genes and CRISPR regions, among other genomic features. The predicted genes were annotated by identifying universal single-copy marker genes (uscMGs) with fetchMGs (v1.2) (Sunagawa et al., 2013), assigning orthologous groups with emapper (v2.0.1) (Huerta-Cepas et al., 2017) based on eggNOG (v5.0) (Huerta-Cepas et al., 2019) and performing queries against the KEGG database (release 2020-02-10) (Kanehisa and Goto, 2000). This last step was performed by aligning the proteins to the KEGG database using DIAMOND (v0.9.30) (Buchfink et al., 2015) with query and subject coverage of ≥70%. The results were further filtered on the basis of the bitscore being ≥50% of the maximum expected bitscore (reference against itself) per the NCBI Prokaryotic Genome Annotation Pipeline (Tatusova et al., 2016). The gene sequences were additionally used as input to identify biosynthetic gene clusters (BGCs) in the genomes using antiSMASH (v5.1.0) (Blin et al., 2019) with default parameters and the different cluster blasts turned on. All genomes and annotations have been compiled along with contextual metadata into the OMD, which is available at https://microbiomics.io/ocean/.

*Gene-level profiling*

Similar to the methods described previously (Salazar et al., 2019; Sunagawa et al., 2015), we clustered the >56.6 M protein-coding genes from the bacterial and archaeal genomes of the OMD at 95% identity and 90% coverage of the shorter gene using CD-HIT (V4.8.1) (Fu et al., 2012) into >17.7 M gene clusters. The longest sequence was selected as the representative gene of each gene cluster. The 1,038 metagenomes were then mapped to the >17.7 M cluster representatives with BWA (*-a*) and the resulting BAM files were filtered to keep only alignments with a percent identity ≥95% and ≥45 bases aligned. Length-normalized gene abundance was calculated by first counting inserts from best unique alignments and then, for ambiguously mapped inserts, adding fractional counts to the respective target genes in proportion to their unique insert abundances.

*Species-level profiling with mOTUs*

The genomes in the extended OMD (augmented with additional MAGs from *Ca.* Eudoremicrobiaceae, see below) were added to the database (v2.5.1) of the metagenomic profiling tool mOTUs (Milanese et al., 2019) to generate an extended mOTUs reference database. Only genomes with at least six out of the 10 uscMGs in single copy were kept (23,528 genomes). The extension of the database resulted in 4,494 additional species-level clusters. Profiling of the 1,038 metagenomes was done using default parameters of mOTUs (v2). Based on the mOTUs profiles, a total of 989 genomes (95% REFs, 5% SAGs and 99.9% belonging to MarDB) contained within 644 mOTUs clusters were not detected. This reflects the various additional marine isolation sources (most of the genomes not detected are associated with organisms isolated from *e.g.*, sediments, marine hosts) of the MarDB genomes. To remain focused on the open ocean environment in this study, we excluded them from downstream analyses if they were not detected or not included in the extended mOTUs database established in this study.

*Clustering and selection of BGCs*

All BGCs from MAGs, SAGs and REFs in the OMD (see above) were combined with the ones identified across all the metagenomic scaffolds (antiSMASH v5.0, default parameters) and processed with BiG-SLICE (v1.1) for feature (PFAM domains) extraction (Kautsar et al., 2021b). On the basis of these features, we computed all-vs-all cosine distances between BGCs and clustered them (average linkage) into gene cluster families (GCFs) and gene cluster clans (GCCs), using a 0.2 and a 0.8 distance threshold, respectively. These thresholds are an adaptation of those previously used with Euclidean distances (Kautsar et al., 2021b) to cosine distances, which alleviate some of the biases of the original BiG-SLICE clustering strategy (Supplemental information).

BGCs were subsequently filtered, keeping only the ones encoded on scaffolds ≥5 kbp and excluding MarDB REFs and SAGs that were not detected in the 1,038 metagenomes (see above). This resulted in a total of 39,055 BGCs encoded by OMD genomes and an additional 14,106 identified on metagenomic fragments (i.e., that were not binned into MAG). These “metagenomic" BGCs were used to estimate the proportion of the ocean microbiome biosynthetic potential not captured by the database (Supplemental information). Each BGC was functionally characterized on the basis of predicted product types as defined by antiSMASH or coarser product classes, as defined in BiG-SCAPE (Navarro-Muñoz et al., 2020). To prevent sampling biases in quantitative analyses (taxonomic and functional compositions of GCCs/GCFs, GCFs and GCCs distances to reference databases as well as GCFs metagenomic abundances), the 39,055 BGCs were further dereplicated by keeping only the longest BGC per GCF per species, leading to a total of 17,689 BGCs.

*Novelty of gene cluster families (GCFs) and clans (GCCs)*

The novelty of GCCs and GCFs were estimated on the basis of distances to databases of computationally predicted (the RefSeq database within BiG-FAM) (Kautsar et al., 2021a) and experimentally validated (MIBIG 2.0) (Kautsar et al., 2020) BGCs. For each of the 17,689 representative BGCs we selected the minimum cosine distance to the respective database. These minimum distances were then averaged (mean) per GCF or GCC as appropriate. A GCF was considered novel if the distance to the database was above 0.2, which corresponds to (on average) the complete separation between the GCF and the reference. For GCCs, we selected 0.4, twice the GCF-defining threshold, to capture remote relationships with the reference.

*Abundance and prevalence of GCFs and GCCs*

The metagenomic abundance of a BGC was estimated as the median abundance of its biosynthetic genes (as defined by antiSMASH), which were available from the gene-level profiles. The metagenomic abundance of each GCF or GCC was subsequently computed as the sum of its representative BGCs (out of the 17,689). These abundance profiles were subsequently cell-normalized using the mOTUs count per sample. The prevalence of a GCF or GCC was computed as the percentage of samples with an abundance >0.

*Structure of the ocean microbiome biosynthetic potential*

Euclidean distances between samples were computed on the basis of the normalized GCF profiles. These distances were dimensionally reduced using UMAP (Becht et al., 2018) and the resulting embedding used for unsupervised density-based clustering with HDBSCAN (McInnes et al., 2017). The optimal minimum number of points of a cluster (and therefore the number of clusters) used by HDBSCAN was determined by maximizing the cumulative cluster membership probability. The identified clusters (as well as random balanced subsamples of these clusters, to account for PERMANOVA’s biases) were tested for significance using PERMANOVA against the non-reduced Euclidean distances. The average genome size of a sample was computed on the basis of relative abundances of mOTUs and the estimated genome sizes of the member genomes. Specifically, the mean genome size for each mOTU was estimated as the mean of the completeness-corrected sizes (e.g., the corrected size of a 75% complete genome with a length of 3 Mbp is 4 Mbp) of its member genomes (after filtering for genomes with a mean completeness of ≥70%). Then, per sample, the average genome size was computed as the sum of the relative abundance-weighted mOTU genome sizes.

*Phylogenomic distribution of BGCs*

The filtered set of BGCs encoded by genomes in the OMD (in scaffolds ≥5 kbp and excluding MarDB REFs and SAGs that were not detected in the 1,038 metagenomes, see above) along with their predicted product classes were displayed on the GTDB bacterial and archaeal trees based on the GTDBTk phylogenomic placement of the genomes (see above). We first reduced the data on a per-species basis, using the genome with most BGCs in that species as a representative. For visualization, the representatives were further binned along the tree and, similarly, for each binned clade the genome containing the most BGCs was selected as representative. BGC-rich species (at least one genome with >15 BGCs) were further analysed by computing the Shannon diversity index of the product types encoded in these BGCs. Chemical hybrids and other complex BGCs (as predicted by antiSMAH) were considered to be from the same product type if all the predicted product types were identical, irrespective of their order within the cluster (e.g., a proteusin;bacteriocin hybrid is identical to bacteriocin;proteusin hybrid).

*Targeted binning of Ca.* Eremiobacterota

After the reconstruction of the first two *Ca*. Eremiobacterota MAGs, we identified six additional ones with ANI >99% (these are included in Figure 3) that were initially filtered out based on contamination estimates (later identified as gene duplications, see below). We additionally recovered bins identified as *Ca*. Eremiobacterota from a different study (Acinas et al., 2019) and used them along with the eight MAGs from our study as a reference for subsampled mapping (5 M reads) of metagenomic reads from 633 eukaryote-enriched (>0.8 µm) samples using BWA (v0.7.17-r1188, *-a* flag). On the basis of enriched specific mappings (after 95% alignment identity and 80% read coverage filtering), 10 metagenomes (expected coverage ≥5X) were selected for assembly and 49 additional metagenomes (expected coverage ≥1X) for abundance correlation. Using the same parameters as described above, these samples were binned and 10 additional *Ca*. Eremiobacterota MAGs were recovered. These 16 MAGs (which excludes the two that were already in the database) bring the total number of genomes in the extended OMD to 34,815. The MAGs were assigned to taxonomic ranks on the basis of their genomic similarity and GTDB placement. The 18 MAGs were dereplicated using dRep into 5 species (within-species ANIs were >99%) and 3 genera (within-genus ANIs ranged between 85-94%) (Barco et al., 2020) within the same family. Species representatives were manually selected on the basis of completeness, contamination and N50. Proposed naming is available in supplementary text (Supplemental information).

*Manual evaluation of Ca.* Eremiobacterota *MAGs*

To evaluate the completeness and contamination of *Ca.* Eremiobacterota MAGs, we assessed the presence of uscMGs, in addition to lineage- domain-specific single-copy marker gene sets used by CheckM and Anvi'o. The identification of duplications among two of the 40 uscMGs was confirmed by phylogenetic reconstruction (see below) to rule out any potential contamination (which would have corresponded to 5% on the basis of these 40 marker genes). Additional inspection of the representative MAGs of the five *Ca.* Eremiobacterota species confirmed low rates of contaminants in these reconstructed genomes on the basis of abundance correlation and sequence composition (Supplemental information) using the Anvi'o interactive interface (Eren et al., 2015).

*Phylogenomics of Ca.* Eudoremicrobiaceae

For phylogenomic analyses, we selected the representative MAGs of the five *Ca*. Eudoremicrobiaceae species, all *Ca*. Eremiobacterota genomes available in GTDB (r89) (Parks et al., 2018) and representatives of additional phyla (incl. UBP13, Armatimonadota, Patescibacteria, Dormibacterota, Chloroflexota, Cyanobacteria, Actinobacteria and Planctomycetota). All these genomes were annotated as previously mentioned to extract single-copy marker genes and to annotate BGCs. GTDB genomes were retained on the basis of the completeness and contamination criteria mentioned above. The phylogenomic analysis was performed using the Anvi'o phylogenomics workflow (Eren et al., 2015). The tree was constructed with IQTREE (v2.0.3) (defaults and *-bb 1000*) (Nguyen et al., 2015) on an alignment (MUSCLE, v3.8.1551) (Edgar, 2004) of 39 concatenated ribosomal proteins identified by Anvi'o, with positions trimmed for coverage in at least 50% of the genomes (Capella-Gutiérrez et al., 2009) and using Planctomycecota as the outgroup based on the GTDB tree topology. Individual trees for the 40 uscMGs were built with the same tools and parameters.

*Trait and lifestyle prediction of Ca.* Eudoremicrobiaceae

We used Traitar (v1.1.2) with default parameters (phenotype, from nucleotides) (Weimann et al., 2016) to predict general microbial traits. We investigated the potential predatory lifestyle on the basis of a previously developed predatory index (Pasternak et al., 2013), which relies on the protein-coding gene content of a genome. More specifically, we used DIAMOND to compare the proteins from a genome to the OrthoMCL database (v4) (Chen et al., 2006) using *--more-sensitive --id 25 --query-cover 70 --subject-cover 70 --top 20* and counted genes that matched predatory and non-predatory marker genes. The index is the difference between the number of predatory and non-predatory markers. As an additional control, we also analysed the genome of *Ca.* Entotheonella factor TSY1 (Wilson et al., 2014) based on its similar characteristics to *Ca.* Eudoremicrobium (large genome size and biosynthetic potential). We further tested a potential link between predatory and non-predatory marker genes with the biosynthetic potential of *Ca*. Eudoremicrobiaceae and found at most one gene (from either type, i.e. predatory/non-predatory, of marker genes) overlapping with BGCs, suggesting that BGCs do not confound the predatory signal. Additional annotations of the genomes to specifically investigate secretion systems, pili and flagella were performed using TXSSCAN (v1.0.2) for unordered replicons (Abby and Rocha, 2017).

*Transcriptomic profiling of Ca.* E. taraoceanii

Transcriptomic profiling was performed by mapping 623 metatranscriptomes from *Tara* Oceans prokaryote- and eukaryote-enriched fractions (Carradec et al., 2018; Salazar et al., 2019; Sunagawa et al., 2020) (with BWA, v0.7.17-r1188, *-a* flag) to the five representative *Ca*. Eudoremicrobiaceae genomes. After 80% read coverage and 95% identity filtering, the BAM files were processed with FeatureCounts (v2.0.1) (Liao et al., 2014) (*featureCounts --primary -O --fraction -t CDS,tRNA -F GTF -g ID -p*) to compute the number of inserts per gene. The resulting profiles were normalized by gene length and mOTU marker gene abundances (median length-normalized insert count of genes with insert count >0) and log2 transformed (Milanese et al., 2019; Salazar et al., 2019) to obtain relative per-cell expression levels of each gene. Such ratios allow for comparative analyses by mitigating the issues of compositionality when working with relative abundance data. Only samples with >5 of the 10 mOTU marker genes were considered for further analyses.

The normalized transcriptomic profiles of *Ca*. E. taraoceanii were dimension-reduced using UMAP and the resulting representation was used for unsupervised clustering using HDBSCAN (see above) to identify expression states. The significance of differences between the identified clusters was tested by PERMANOVA in the original (non-reduced) distance space. Differential expression between these states was tested across 201 KEGG pathways identified in the genome (see above) and 6 functional groups, namely: BGCs, secretion systems and flagellar genes from TXSSCAN, degradative enzymes (proteases and peptidases) from prokka and predatory and non-predatory markers from the predatory index. Per sample, we computed the median normalized expression for each category (note that BGC expression was itself computed as the median expression of the biosynthetic genes of that BGC) and tested for significance (FDR-corrected Kruskal Wallis test) across the different states.

*Materials for the expression of a Ca.* E. malaspinii-*specific bacteriocin cluster (BGC 75.1)*

Synthetic genes were purchased from GenScript and PCR primers were ordered from Microsynth AG (Switzerland). Phusion polymerase from Thermo Fisher was utilized for DNA amplification. NucleoSpin plasmid and NucleoSpin Gel and PCR Clean-up kits from Macherey-Nagel were used to purify DNA. Restriction enzymes and T4 DNA ligase were purchased from New England Biolabs. Chemicals were purchased from Sigma Aldrich, with the exception of isopropyl-β-D-1-thiogalactopyranoside (IPTG) (Biosynth AG) and 1,4-dithiothreitol (DTT, AppliChem) and utilized without further purification. The antibiotics chloramphenicol (Cm), spectinomycin dihydrochloride (Sm), ampicillin (Amp), gentamicin

(Gt) and carbenicillin (Cbn) were purchased from AppliChem. The media components Bacto Tryptone and Bacto Yeast Extract were purchased from BD Biosciences. Sequencing grade trypsin was purchased from Promega AG.

*Cloning of embA, embM and orf3 (embI) for protein expression*

The genes *embA* (locus: MALA_SAMN05422137_METAG-scaffold_127-gene_5), *embM* (locus: MALA_SAMN05422137_METAG-scaffold_127-gene_4), *embAM* (including intergenic region) were ordered as synthetic constructs in pUC57(Amp$^R$), with and without codon-optimization for expression in *E. coli*. *embA* was subcloned in the first multiple cloning site (MCS1) of pACYCDuet-1(Cm$^R$) and pCDFDuet-1(Sm$^R$) with BamHI and HindIII cut sites. *embM* and *embMopt* (codon optimized) were subcloned in MCS1 of pCDFDuet-1(Sm$^R$) with BamHI and HindIII and in the second multiple cloning site (MCS2) of pCDFDuet-1(Sm$^R$) and of pRSFDuet-1(Kan$^R$) with NdeI/XhoI. *embAM* was subcloned in pCDFDuet1(Sm$^R$) with BamHI and HindIII cut sites. The gene *orf3/embI* (locus: MALA_SAMN05422137_METAG-scaffold_127-gene_3) was constructed by overlap extension PCR with primers EmbI_OE_F_NdeI and EmbI_OE_R_XhoI, digested with NdeI/XhoI and ligated to pCDFDuet-1-EmbM(MCS1), which was digested with the same restriction enzymes (Table S5). Restriction enzyme digestions and ligations were performed following the New England Biolabs manufacturer procedures.

All constructs generated above were transformed in chemically-competent *E. coli* DH5α and plated on LB agar with appropriate antibiotic selection. Plasmids were purified from single colonies and sequenced using sequencing primers to verify proper insertion of genes (Table S5). *embA* and *embAM* were additionally subcloned in a modified pLMB509m(Gt$^R$) vector for *Microvirgula aerodenitrificans* expression via Gibson assembly, with the inclusion of an N-terminal His$_6$ purification tag in the final EmbA protein product (Bhushan et al., 2019; Lotti and Piel). Gibson assembly primers EmbA_F_plmb, EmbA_R_plmb, Plmb_F_EmbA, Plmb_R_EmbA, are listed in Table S5. Transformation in *E. coli* DH5α, isolation of plasmids and validation of correct clones by sequencing was followed by transformation of NHis$_6$-EmbA-pLMB509m and NHis$_6$-EmbAM-pLMB509m into *E. coli* SM10 for conjugation.

*Heterologous expression and purification for protein isolation for in vitro assays*

Chemically-competent *E. coli* BL21(DE3) were transformed with pCDFDuet-1-EmbA(MCS1) and pCDFDuet-1-EmbM(MCS1). The same conditions were used for expression of both N-terminally His$_6$-tagged proteins. Overnight cultures were prepared from single colonies and used to inoculate (1% v/v) TB media (2x200 mL) in 1-L baffled Erlenmeyer flasks supplemented with spectinomycin (50 mg/mL). Cells were grown at 37°C, 200 rpm, until $OD_{600\ nm}$ ~ 1.0, cooled on an ice bath for 20 min, and induced with a final concentration of 0.5 mM IPTG. The cultures were further incubated at 16°C, 180 rpm for 18-20 h, and subsequently harvested by centrifugation (8,000×g, 20 min) and frozen.

Purifications of NHis$_6$-EmbA and NHis$_6$-EmbM were carried out at 4°C, utilizing the same procedure for both proteins. Cell pellets were resuspended in 5 mL/g of lysis buffer (50 mM

Tris, 300 mM NaCl, 5 mM imidazole, 10% glycerol, pH 7.8). The suspension was supplemented with lysozyme (1 mg/mL), DNase I (10 U/mL) and protease inhibitor cocktail (0.2% v/v) and stirred at 37°C for 30 min. After cooling the suspension for 15 min on an ice bath, cells were lysed by sonication (30% amplitude, 10s on/off cycles, for a total of 3 minutes), and the clarified lysate was obtained by centrifugation (27,000×g, 30 min). The supernatant was loaded on 4 mL of Ni-NTA agarose resin that had been equilibrated with a lysis buffer in a fritted purification column. The resin was washed with 10 column volumes (CV) of lysis buffer, 3 CV of wash buffer (50 mM Tris, 300 mM NaCl, 40 mM imidazole, 10% glycerol, pH 7.8) and finally eluted with 3 CV of elution buffer (50 mM Tris, 300 mM NaCl, 250 mM imidazole, 10% glycerol, pH 7.8), in 1.5 mL fractions. Elution fractions were analyzed by SDS-PAGE, pooled, and concentrated in spin concentrators with appropriate molecular weight cutoff. $NHis_6$-EmbA and $NHis_6$-EmbM were buffer-exchanged using a PD MiniTrap G25 column pre-equilibrated with G25 buffer (50 mM Tris, 300 mM NaCl, 10% glycerol, pH 8.0). The concentration of buffer-exchanged proteins was determined by measuring the absorbance of purified proteins at 280 nm and using the calculated values for molecular weight and extinction coefficient for each protein.

*In vitro enzymatic activity assays with EmbA and EmbM*

Extensive screening of enzymatic reaction parameters, including temperature, time, enzyme and substrate concentration, buffer pH and salinity resulted in the following best condition set for EmbM turnover. EmbA was added to a glass vial to a final concentration of 200 mM. Final concentrations of 2 mM of $MgCl_2$ and 2 mM of adenosine 5'-triphosphate (ATP) were added to the reactions. EmbM was not added to control reactions and added to a final concentration of 10 mM in turnover experiments. The enzymatic reaction was stirred at 37°C for 72 h, and 2 mM of ATP were supplemented to the reaction mixture every 24 h. Reaction scales ranged from 100 µL, for analytical purposes, to 3 mL for product isolation.

Modified EmbA was proteolyzed with trypsin for mass spectrometry analysis and with LahT150 for mass spectrometry analysis and product isolation (Bobeica et al., 2019). Trypsin cleavage was performed by diluting the reaction mixture with 2x trypsin buffer (50 mM Tris, 5 mM $CaCl_2$, pH 8.0), adding 1:20 trypsin : EmbA and incubating overnight at 37°C. LahT150 cleavage was performed by adding LahT150 in a 1:10 ratio to EmbA, and incubating at room temperature overnight for small scale reactions or for 24 h, for reactions larger than 1 mL.

Large scale enzymatic reactions were purified using solid-phase extraction (SPE) with a Phenomenex Strata C18-E reverse phase column (2g sorbent). The sorbent was first washed with 24 mL MeOH and equilibrated with 24 mL $H_2O$ (+0.1% formic acid). The proteolysis reactions were loaded onto the sorbent, which was then washed with 24 mL $H_2O$ (+0.1% formic acid). The peptide products were eluted with 24 mL of 1:1 MeCN : $H_2O$ (+0.1% formic acid) and 12 mL MeCN (+0.1% formic acid). Elution fractions were pooled, dried on a Genevac concentrator, and stored at -20°C.

*Co-expression of EmbA, EmbM and Orf3 (EmbI) in heterologous hosts and purification*

A wide variety of *E. coli* co-expression conditions of EmbA, EmbM and Orf3 were screened (see Table S5). In general, chemically competent BL21(DE3) or Tuner(DE3) were transformed with different combinations of the constructs described above, and selected with appropriate antibiotics on LB plates. Overnight cultures were prepared from single colonies and 1% v/v of culture was utilized to inoculate 200 mL of media (TB, LB, XPPM) (Bode et al., 2019) supplemented with appropriate antibiotic selection in 1-L baffled Erlenmeyer flasks. Cells were grown at 37°C, 200 rpm until $OD_{600\ nm}$ ~1.2. For low temperature growths, cultures were subsequently cooled in ice baths for 20 min, induced with 0.5 mM IPTG and incubated at 16°C, 180 rpm. Incubation times varied from 24 h to 7 days. High temperature growths were induced with a final concentration of 0.5 mM IPTG without cooling and incubated at 37°C, 200 rpm from 6 h to 16 h. Cultures were harvested by centrifugation (8,000×g, 20 min), frozen, and purified as described above. Proteolysis reactions with trypsin or LahT150 for mass spectrometry analysis were performed as discussed in the previous section.

Transformation of *M. aerodenitrificans* DSMZ 15089 with $NHis_6$-EmbA-pLMB509m and $NHis_6$-EmbAM-pLMB509m was achieved following published procedure (Bhushan et al., 2019). Culturing conditions followed an adaptation of a previously described method (Bhushan et al., 2019; Lotti and Piel). Cultures were harvested, stored and purified as described above. Trypsin digestion was used for mass spectrometry analysis.

*β-Elimination of phosphorylated peptides*

Phosphorylated peptide intermediates obtained via co-expression of EmbA and EmbM were submitted to β-elimination conditions in 100 μL scale. EmbA (200 μM) in G25 buffer was used either as an intact protein or as a trypsin digest product. The solution's pH was adjusted to pH 13 with 1 M NaOH, and the elimination reaction proceeded at 37°C for 4 h to afford dehydrobutyrine (Dhb)-containing products. Derivatization of Dhb was performed by adding a final concentration of 50 mM DTT to the reaction. The pH of the solution was adjusted to 7 with HCl (aq.) prior to mass spectrometry analysis.

β-Elimination of phosphorylated peptide intermediates obtained through *in vitro* enzymatic reactions was carried out with EmbA which had been proteolyzed with LahT150 and purified via SPE as described above. EmbA (0.6 mmol) was resuspended in 3 mL of $H_2O$, and the pH of the solution was adjusted to 14 with 1 M NaOH. The reaction stirred at 37°C for 48 h, and was subsequently neutralized to pH 7 with HCl (aq.). SPE purification was carried out as described above.

*HPLC-HR-MS and MS/MS Analysis*

HPLC-HR-MS and MS/MS analyses were performed on a Thermo Scientific Dionex UltiMate 3000 UHPLC coupled to a Thermo Scientific Q Exactive mass spectrometer using heated electrospray ionization in positive ion mode with a Phenomenex Kinetex 2.6 μm XB-C18 100 Å (150×4.6 mm) column. The column temperature was set to 50°C and the flow rate to 0.5

mL/min. Samples were centrifuged before injections and target peptides were eluted with a gradient of 15-55% MeCN (+0.1% formic acid) over 15 min. Full MS was performed at a resolution of 70,000 (AGC target 1e6, maximum IT 100 ms) and parallel reaction monitoring (PRM) was done at a resolution of 17,500 (AGC target 2e5, maximum IT 100 ms, isolation windows in the range of 2.0 *m/z*).

*Bioactivity assays*

*Escherichia coli* DSM 1103, *Staphylococcus aureus subsp. aureus* ATCC 29213, *Pseudomonas aeruginosa* DSM 1117, *Acinetobacter baumanii* DSM 30007, *Enterococcus faecalis* DSM 2570, *Rhodococcus* sp. L233*, Aquimarina sp. Aq135, Rheinheimera aquimaris* B26*, Vibrio spartinae* (salt marsh isolate), *Pseudoalteromonas rubra* DSM 6842, *Saccharomyces cerevisiae* W301-1A and *Pichia pastoris* (*Komagataella phaffii*) NRR Y-11430 were tested for antimicrobial activity with the dehydrated peptide from BGC 75.1. Bioactivity assays were carried out in accordance with the 2003 guidelines of the Clinical and Laboratory Standards Institute (CLSI) using the microtiter method.

*E. coli, S. aureus, A. baumanii,* and *P. aeruginosa* overnight cultures were grown in LB at 37°C. *P. rubra, R. aquimaris* (37°C)*, V. spartinae* (30°C), *P. rubra and Aquimarina* sp. Aq135 (24°C) were cultured in marine broth at their respective growth temperature optimums indicated in parentheses. *Rhodococcus* sp. L233 was cultured in R2A medium at 24°C. *S. cerevisiae* and *P. pastoris* were cultured in YPD medium at 28°C. An overview of strains, growth conditions, and taxonomy is provided in Table S5.

Microbial seed cultures were initiated by inoculating 5 mL of medium for each strain and by incubating overnight shaking at 200 rpm. Each culture was then diluted with their respective growth media to an initial $OD_{600\ nm}$ of 0.02 in 80 µL volume per well in sterile 96-well plates (one per strain tested). Assays were set-up in duplicates, with appropriate controls including solvent (water) controls and positive controls consisting of two different broad-spectrum antibiotics (chloramphenicol and ampicillin) with 50 µM final concentrations. Cycloheximide and benomyl were used as positive controls for *S. cerevisiae* and *P. pastoris*.

Two final concentrations of the peptide, using water as a solvent, were tested: 50 and 25 µM. 96-well plates were parafilmed and incubated at room temperature without shaking. $OD_{600\ nm}$ was determined after 10s of plate agitation at the following time points: 2h, 4h, 6h, 8h, 18h, 21h, 24h and 48h.

### Supplemental Information

Supplemental information for this manuscript includes:

- Supplementary Text
- Figures S1 to S16
- Captions for Tables S1 to S5

Other Supplementary Materials for this manuscript include the following:

- Tables S1 to S5

### Supplementary Text

#### Improved reconstruction of metagenome-assembled genomes (MAGs)

*Abundance correlation improves the number and quality of recovered MAGs*

To evaluate the overall impact of abundance correlation on the recovery of MAGs, we reconstructed MAGs with and without abundance correlation in a random subset of metagenomes (20 from *Tara* Oceans virus- and prokaryote-enriched datasets, five from Malaspina, 10 from bioGEOTRACES and five from each time-series study). We computed how well a sample was binned into MAGs using the sum of the quality scores (Q') (Methods) of the reconstructed MAGs (*i.e.*, after filtering incomplete or contaminated bins) to capture variations in both the number and quality of the recovered MAGs. For each sample, we then computed the ratio of this metric between binning with and without abundance correlation (Figure S2). We found the ratios to be >1 (except for a single virus-enriched sample from which no MAG could be recovered with either strategies), with a median ratio of 2.3 across all randomly selected samples. This increase is due to both an increased number (mean 2.7 times) and improved quality (mean +20%) of the recovered MAGs, indicating the use of abundance correlation to be strictly beneficial. Out of the 26,293 MAGs reconstructed in this study, we estimated that more than 95% benefited from abundance correlation, as they were detected in ≥3 samples (75% in ≥10 samples).

*Reconstructed MAGs have improved qualities compared to previous efforts*

We compared the quality of MAGs reconstructed in this study to other efforts of reconstructing ocean microbial genomes. Comparisons were performed on the basis of the quality scores (Q') described above (Methods).

The comparison to two datasets of external MAGs reconstructed with automated workflows (Parks et al., 2017; Tully et al., 2018) was performed based on shared GTDB (Parks et al., 2018) species-level annotations (i.e. 95% ANI single-linkage). An additional comparison was made with manually-curated MAGs (Delmont et al., 2018) included in the OMD on the basis of MAGs sharing the same species-level cluster (95% ANI clustering). In both comparisons, samples that were binned in this study but not included in the publicly available MAGs datasets were excluded. For all the species that had at least one MAG reconstructed in this study and at least one from either dataset, we calculated the difference in Q' score of the best-scoring MAGs.

The results revealed that the approach of combining single-sample assemblies with large-scale abundance correlations achieved on average significantly higher community-defined quality scores (Bowers et al., 2017) than automatically generated MAGs (Parks et al., 2017; Tully et al., 2018), and even manually-curated, co-assembled MAGs (Delmont et al., 2018) (Figure S8).

*Enhanced recovery of mobile genetic elements*

We further sought to investigate the ability of the binning strategy to recover mobile genetic elements (MGEs) within MAGs, since this has been reported as a challenge associated with MAGs reconstruction (Maguire et al., 2020). More specifically, we focused on plasmids and phages, with the expectation that recovering plasmids would be desirable while associating virus fragments (except for prophages) to MAGs could be considered as contamination. To this end, all the ≥80M metagenomic scaffolds were annotated with PlasmidFinder (v2.1, with *-l 0.66 -t 0.5*) (Carattoli et al., 2014), PlasFlow (v1.1.0, with *--threshold 0.7 --batch_size 10000*) (Krawczyk et al., 2018), cBar (v1.2) (Zhou and Xu, 2010), VirSorter (v1.0.5, with *--db 2 --no_c*) (Roux et al., 2015), DeepVirFinder (v1.0) (Ren et al., 2020), EukRep (v0.6.6, with *--min 1000 --seq_names -m balanced --tie skip*) (West et al., 2018) and ccontigs (https://github.com/Microbiology/ccontigs/). These tools allowed the identification of plasmids (first three tools) and viruses (VirSorter and DeepVirFinder) in the metagenomes, while accounting for eukaryotic scaffolds (using EukRep) that can be detected as false positives, and complete circular molecules (ccontigs).

We found plasmids to be binned within MAGs at a much improved rate (Figure S9) compared to previous reports (Maguire et al., 2020). Specifically, the difference in probability of binning a plasmid vs a chromosomal fragment (biased against plasmids) ranged from -10 to -17%, a leap compared to previously reported difference of at best -50%. Furthermore, the rates of viral fragments (excluding prophages) associated with MAGs below 0.1% suggest little sensitivity of our reconstruction method to viral contaminants, which would not be picked up by usual quality metrics relying on prokaryotic marker genes.

*Evaluation of chimera through taxonomic uniformity*

A particularly critical issue with the reconstruction of MAGs is that the reliance on single-copy marker genes counts for contamination estimation can still lead to chimerism, with populations sometimes from completely different clades being mixed within a single genome (Chen et al., 2020; Shaiber and Eren, 2019). To quantify this risk for the genomes in the database, we annotated 10 single-copy marker genes (Milanese et al., 2019) and evaluated the homogeneity of their taxonomic annotation for each genome using STAG (v0.7, https://github.com/zellerlab/stag). Briefly, STAG annotates marker gene sequences along a provided taxonomic tree, here the NCBI taxonomy available from the progenome database (v2) (Mende et al., 2020), using a LASSO classification model at each bifurcation of the tree. We evaluated, for each genome, the congruence of these annotations, defining the following categories: "No annotation" if a maximum of one gene was annotated; “Agreeing" if all genes had the same annotation; "Majority agreeing" if more than half the genes had the same annotation and “Not agreeing” otherwise. Notably, across all MAGs the rate of disagreement was <1% with that rate being ~0.1% for MAGs with differential coverage index ≥10 (i.e. 75% of the MAGs) (Figure S10).

#### Biosynthetic potential of the ocean microbiome

*Improved BGC clustering and BGC class enrichment in the ocean*

The recent analysis of ~1.2 M BGCs predicted from ~190,000 genomes deposited at RefSeq relied on pairwise euclidean distances between BGCs based on BGC features followed by BIRCH clustering to identify GCFs (Kautsar et al., 2021a). Among the identified GCFs, nearly 40% of them were predicted to encode for NRPS and only ≤5% were predicted to encode for RiPPs. Applying the same approach to the set of 39,055 filtered BGCs in the OMD, we found, by contrast, that most GCFs were predicted for other classes of natural products, followed by terpenes. Notably, 8% of the GCFs were predicted to encode RiPPs (Table S2).

However, this previous clustering approach was found to have lower sensibility for product classes usually encoded by fewer features, e.g., terpenes and RiPPs (Kautsar et al., 2021b). To address that issue, we adapted the previous strategy by using cosine rather than euclidean distances and average linkage rather than BIRCH clustering. Comparing both approaches to the set of 39,055 filtered BGCs in the OMD, we found the number of GCFs to increase from 5,195 to 6,907 and, most strikingly, the proportion of RiPPs among these was increased by over two-folds (from 8 to 17%). The proportion of terpenes also grew from 25 to 32%, while, relatively, other classes had their proportions reduced (Table S2).

*The OMD provides genomic context to most BGCs*

In addition to the 39,055 BGCs identified in the genomes of the database, we found 14,106 that were encoded on metagenomic fragments that were not binned into MAGs. To evaluate how much of the biosynthetic potential of the ocean microbiome was not captured by the OMD, we grouped both sets of BGCs and clustered them into GCCs and GCFs (Methods). We found that 95% and 81% of these GCCs and GCFs, respectively, had at least one representative encoded in a genome within the OMD. Notably, we found that BGCs encoding for a specific product type, nucleosides, were poorly recovered by MAGs. Instead, these BGCs were particularly enriched in predicted phages (Figure S11), suggesting that they may carry BGCs usually involved in producing hypermodified nucleosides. These clusters span 87 different GCFs mostly from two GCCs. Although phages are known to encode and use DNA hypermodifications (Lee et al., 2018), this finding suggests such modifications may be widespread in marine phages and possibly linked to an arms race with bacteria defense mechanisms (Weigele and Raleigh, 2016).

*The ocean microbiome specialized metabolism*

We found several GCCs to harbour a particularly large phylogenomic distribution (>10 phyla) and to be particularly prevalent in the metagenomic dataset (detected in >80% of the samples). Interestingly, these were predicted, for example, to encode for heterocyst-like glycolipid (hglE-KS), polyunsaturated fatty acids (PUFA), aryl-polyenes, ectoines and siderophores which can be involved in membrane fluidity, oxidative and osmotic stress resistance as well as iron uptake respectively (Allemann et al., 2019; Moore et al., 2013; Pastor et al., 2010; Sutherland et al., 2020). Together, this suggests that these GCCs could capture an ocean specialized metabolism, that reflects microbial adaptation to the marine environment, rather than a lineage-specific secondary metabolism.

*Machine learning-based (GECCO) detection of potential BGCs*

We additionally complemented the rule-based BGC prediction approach used by antiSMASH with GECCO (v0.4.4) (https://gecco.embl.de), a recently developed, machine learning-based approach, which has the potential to detect BGCs of unknown architecture. Notably, when applied to the same set of genomes, GECCO (using its default parameters) predicted a total of 357,541 clusters, as opposed to 51,851 predicted by antiSMASH (this number is larger than the reported 39,045 BGCs in the main text, as it includes scaffolds <5 kbp and MarDB genomes that were not detected across the 1,038 metagenomes). Although this seven-fold increase likely includes false positives, this reserve of candidate BGCs may also include novel classes of natural products that are not picked up by usual prediction approaches (Supporting data).

#### The marine lineage of *Ca.* Eremiobacterota

*Binomial naming of a new marine Ca.* Eremiobacterota *lineage*

Based on whole genome ANIs, taxonomic annotations and phylogenomic analyses (Figure 4) (Methods), we identified five species from three genera belonging to the same family. We propose the following names (Murray et al., 2020):

‘*Candidatus* Eudoremicrobium’ (Eu.do.re.mi.cro'bi.um; N.L. fem. n. Eudore, the Nereid, sea deities in Greek mythology, of fine gifts from the sea; N.L. neut. n. microbium, a microbe; N.L. neut. n. Eudoremicrobium, a gifted microbe from the sea);

‘*Candidatus* Eudoremicrobium malaspinii’ (malaspinii; after Malaspina, the name of the expedition that recovered the genetic material of the microbe). Type species of the genus with the genome designated as MALA_SAMN05422137_METAG_HLLJDLBE as type material;

‘*Candidatus* Eudoremicrobium taraoceanii’ (taraoceanii; after *Tara* Oceans, the name of the expedition that recovered the genetic material of the microbe). With the genome designated as TARA_SAMEA2623601_METAG_PIAMPJPB as type material;

‘*Candidatus* Amphithoemicrobium’ (Am.phi.tho’e.mi.cro'bi.um; N.L. fem. n. Amphithoe, the Nereid, sea deities in Greek mythology, she that flows around; N.L. neut. n. microbium, a microbe; N.L. neut. n. Amphithoemicrobium, a microbe from the sea flowing around, referring to the original observation of its distribution across the oceanic depth layers);

‘*Candidatus* Amphithoemicrobium indianii’ (indianii; after the Indian Ocean, basin where the genetic material of the microbe was recovered). With the genome designated as TARA_SAMEA2730749_METAG_OLPPLKCL as type material;

*‘Candidatus* Amphithoemicrobium mesopelagicum (mesopelagicum; after the mesopelagic depth layer, where the microbe was originally observed to be most abundant). With the genomes designated as TARA_SAMEA2623054_METAG_OCMKBGHM as type material;

*‘Candidatus* Autonoemicrobium’ (Au.to.no’e.mi.cro'bi.um; N.L. fem. n. Autonoe, the Nereid, sea deities in Greek mythology, with her own mind; N.L. neut. n. microbium, a microbe; N.L. neut. n. Autonoemicrobium, a microbe from the sea with its own mind, based on the original recovery of a single species in the genus);

*‘Candidatus* Autonoemicrobium septentrionale’ (septentrionale; from septentrio, the north, referring to the original detection of the microbe almost exclusively in the northern hemisphere). With the genomes designated as TARA_SAMEA2623601_METAG_LGBFILLL as type material;

On the basis of these clades, we further propose *‘Candidatus* Eudoremicrobiaceae’ (fam. nov.), *‘Candidatus* Eudoremicrobiales’ (ord. nov.) and *‘Candidatus* Eudoremicrobiia’ (class nov.);

*Manual inspection of the species representatives*

We used Anvi'o to manually inspect the abundance correlation patterns of each of the representative genomes (Figure S12). The uniform read coverage across the scaffolds and stable GC content support a high quality of the identified MAGs. We did note some irregularities in the coverage, yet careful inspection of one such region with higher coverage (e.g., Figure S12A) revealed that this fraction of a larger scaffold was flanked by recombinases, indicating that it is probably present in several copies within the genome, but collapsed in the present assembly. We additionally tested the quality of the *Ca*. E. malaspinii representatives by inspecting the assembly graphs. Briefly, we extracted reads mapping to the genome from several samples and conducted a specific assembly (same parameters as before) to check whether scaffolds would be connected in the assembly graph, suggesting that connected fragments could be part of a single chromosome (Figure S12B). We indeed found that over 99% of the genome was connected to each other, including through what appears to be complex repeat regions.

*Refining the contamination estimates of Ca.* Eudoremicrobiaceae *MAGs*

Initial contamination estimates based on universal single-copy marker genes (CheckM, Anvi'o) were initially above the recommended 10% for some of the recovered MAGs. However this is not unexpected since considering that this set of genes is not optimized for such underexplored phyla (e.g., CheckM later integrated an alternative set of markers more appropriate for CPR genomes) and among the markers used by CheckM and Anvi'o, several are found in more than 1.1 average copies (Olm et al., 2020). We therefore additionally used the set of 40 uscMGs that were selected with more stringent parameters. Indeed, based on these 40 markers the contamination estimates of these MAGs ranged from 2.5 to 5% with COG0124 duplicated in the five species and COG0522 in three of them. To test whether these duplications were due to actual contamination or biological duplication events, we constructed phylogenetic trees based on these genes (Methods). We found, for instance, COG0124 to be consistently duplicated across the MAGs reconstructed in our study as well as external MAGs belonging to related *Ca*. Eremiobacterota lineages (Figure S13). Interestingly, one copy of COG0124 displayed a pattern similar to the phylogenomic analyses while the other was more closely related to Actinobacteria and Planctomycetota. These results support a duplication of the marker gene through introgression rather than contamination during the reconstruction process. Interestingly, these potential introgressions for BGC-rich phyla could be linked to the increased genome sizes and biosynthetic potential observed within the new species.

#### *Ca*. Eudoremicrobiaceae secondary metabolism

*A shared biosynthetic potential*

The biosynthetic potential of *Ca*. Eudoremicrobiaceae (antiSMASH predictions) was clustered using BiG-SLICE (v1.1.0). This clustering was manually curated to identify core and clade specific BGCs. On that basis, we identified six candidate BGCs shared across the five *Ca*. Eudoremicrobiaceae species that constitute the shared biosynthetic potential of the lineage. However, the bacteriocin DUF692 was present in four out of five species, only missing in *Ca*. A. mesopelagicum. This species was only represented by a single MAG with a completeness estimate of 94.3% and using this metric as a probability estimate to find a feature in the reconstructed genome, there is a 5.7% chance that this cluster would be missing by chance. Considering that this probability wouldn't meet significance criteria as well as the phylogenetic relationship between the five species, we conclude that this cluster is most likely shared by the five species.

Interestingly, some of these clusters have similarity with characterized biosynthetic gene clusters from the curated MIBiG database (Kautsar et al., 2020) providing insights into the potential chemical and functional profiles of *Ca*. Eudoremicrobiaceae secondary metabolites.

Siderophore

A classical siderophore biosynthesis cluster conserved across all *Ca*. Eudoremicrobiaceae species which likely has a role in iron scavenging that may provide a fitness advantage in iron-limited marine regions or link to their putative predatory behavior (Moore et al., 2013).

Ectoine

Another conserved gene cluster encoding an ectoine synthetase suggests that *Ca*. Eudoremicrobiaceae spp. may produce the osmolyte ectoine. Ectoine and related products have been shown to help microorganisms survive osmotic stress such as fluctuating salinity conditions in variable ocean environments (Pastor et al., 2010; Widderich et al., 2014).

*Type III polyketide synthase*

A type III polyketide synthase cluster conserved across all *Ca*. Eudoremicrobiaceae species shared similarity with two of the three genes of the characterized BGCs encoding for alkylpyrone or alkylresorcinol-type metabolites. These metabolites typically have long aliphatic tails that are believed to incorporate into cytoplasmic membranes and play a role in regulating membrane rigidity (Funa et al., 2006; Funabashi et al., 2010). *Ca*. Eudoremicrobiaceae MAGs were recovered from different ocean fractions with estimated particle sizes ranging between 0.8 and 20 µm, suggesting that they may form microcellular aggregates such as biofilms. Biofilm formation can also be regulated by secondary metabolites, such as alkylpyrone-type molecules. For example, in *Bacillus* spp., exogenous addition of 4-hydroxyl alkylpyrones resulted in a hyper-wrinkled biofilm morphology (Grubbs et al., 2017).

Polyunsaturated fatty acids and hydrocarbons

A polyketide cluster conserved in the *Ca*. Eudoremicrobiaceae family encodes enzymes with similarity to the multimodular polyunsaturated fatty acid synthase known to produce long-chain polyunsaturated fatty acids (PUFAs). Interestingly, PUFA genes are co-localized with *oleBCD* genes known to form a complex for the production of long-chain ($C_{31+}$) polyunsaturated hydrocarbons (Allemann et al., 2019). These long-chain hydrocarbons likely alter membrane fluidity in response to variable temperature and pressure conditions which *Ca*. Eudoremicrobiaceae members are likely to experience in the marine environment. Metatranscriptomic analysis suggested the expression of *Ca*. Eudoremicrobiaceae polyunsaturated hydrocarbon biosynthetic cluster is constitutive. This finding is consistent with previous studies that have shown that transcription of PUFA genes in the deep-sea bacterium *Photobacterium profundum* strain SS9 does not change under variable cultivation conditions such as increased pressure or reduced temperature despite an observed increase in PUFA production (Allen and Bartlett, 2002). An independent study found that hydrocarbons produced via the *oleABCD* pathway are also constitutively produced (Sukovich et al., 2010).

*Ca.* Eudoremicrobium-*specific biosynthetic potential*

T1PKS/3*NRPS

An interesting BGC conserved exclusively within the *Ca*. Eudoremicrobium genus encodes a hybrid T1PKS/NRPS megasynthase with two neighboring NRPS modules forming an interleaved cluster 77 kb in length (Figure 4D). The total of four adenylation domains in the cluster were all predicted to have specificity for aromatic amino acids such as phenylalanine, tryptophan, hydroxyphenylglycine or dihydroxybenzoate by NRPSsp (Prieto et al., 2011). The BGC shares some similarities with the bacilysin pathway found in *Bacillus* spp. (Parker and Walsh, 2013) Bacilysin is a dipeptide ‘Trojan horse’ antibiotic in which L-alanine is bound to L-anticapsin for export prior to peptidase cleavage and release of the active anticapsin drug. Both pathways encode relatively small peptidic products with aromatic side chains that are highly modified by an abundance of reductases and dehydrogenases in the cluster (Figure S14A). However, unlike the bacilysin pathway which produces a dipeptide, we predict the *Ca*. Eudoremicrobiaceae BGC encodes a hybrid PKS/NRPS final product, which is likely also glycosylated as suggested by the presence of two glycosyltransferases encoded in the cluster. Additional Fe(II)/α-ketoglutarate-dependent oxygenases and

*O*-demethylase Rieske oxygenases suggests the final product is likely more oxidized than bacilysin.

Candidate proteusin 54.1

The most complex, lineage-specific RiPP BGC identified in *Ca*. Eudoremicrobiaceae harbors a number of varied tailoring enzymes and likely yields a highly modified peptide product (Figure S14B). The precursor peptide contains a Nif11-type leader portion (as opposed to the usual NHLP leader of proteusins), with a canonical C-terminal Gly-Gly cleavage motif. Notably, a second Gly-Gly site is identified within the predicted core region. A LanM-type lanthionine synthetase (encoded by *orf9*) putatively installs up to four lanthionine bridges on the precursor peptide, as four cysteine residues are present in the core peptide. Predicted radical SAM (rSAM) epimerases (*orf10, orf11, orf19*) homologous to PoyD and OspD are likely to install D-amino acids on the peptide, and $B_{12}$-dependent radical SAM enzymes (*orf12*, *orf21*) homologous to C-methyltransferases potentially methylate carbons in the precursor (Freeman et al., 2012; Morinaka et al., 2014). Although precursor peptides are highly divergent, the presence of D-amino acids and (methyl)lanthionine rings in the product of cluster 54 suggests some similarities to the structure of landornamides (Bösch et al., 2020). A number of other metalloenzymes belonging to the rSAM, P450, mononuclear non-heme iron α-ketoglutarate dependent families, encoded by *orf13*, *orf14*, *orf16*, *orf17*, *orf20*, likely oxidize the peptide product. Finally, predicted transporters (*orf3*-*orf7*), including some with an N-terminal C39 peptidase, are expected to both cleave the Nif11-type leader peptide and export the mature natural product to the extracellular milieu. This cluster is also co-localized with an uncharacterized family of phage plasmid transfer proteins (encoded by *orf22*) found in the plasmid SCP1 of *Streptomyces coelicolor* and various *Mycobacterium* phage genomes, suggestive of cluster mobility.

Proteusin cluster 34.1

This proteusin cluster contains a precursor peptide rich in small hydrophobic amino acids and harboring an $NX_5N$ pattern of residues in the core region. Together with the presence of other key maturases such as rSAM epimerases (Figure S14C), this cluster is highly characteristic of pore-forming β-helix peptides, such as the highly cytotoxic polytheonamides and aeronamides (Bhushan et al., 2019; Freeman et al., 2012; Hamada et al., 2005)

*Deep-sea specific BGCs*

Aryl polyene

The deep-sea *Ca*. Eudoremicrobium (*Ca*. E malaspinii) also have a unique aryl polyene cluster, which is surprising given that aryl polyenes typically play a role in protection from photodamage by visible light by quenching reactive oxygen species (ROS). However, visible light does not reach depths of 2,000 – 4,000 m where organisms with the aryl polyene cluster are exclusively found. This suggests the aryl polyene product might play a different ecological role, or that ROS may be generated from other sources. A recent study found that marine microbial ROS production through one-electron reduction of $O_2$ to superoxide production plays a larger role in the marine oxygen cycle than previously realized

(Sutherland et al., 2020). Indeed, 'dark' biological superoxide production was demonstrated in most major groups of marine microbes. This suggests ROS mitigation strategies such as through aryl polyene production may also be essential for the survival of bathypelagic microbes even in the absence of visible light.

*Bacteriocin cluster 75.1*
Additionally, we found the deep ocean species (*Ca*. E. malaspinii) representatives to encode for a unique bacteriocin cluster within these clades. Due to this particularity, we selected this cluster for further characterization.

*Experimental validation of Ca.* E. malaspinii *Bacteriocin cluster 75.1*

*In silico* cluster analysis
Protein sequence similarity (Johnson et al., 2008) shows EmbA harbors an N-terminal leader region commonly found in ribosomal natural products, with homology to the Nif11 enzyme, involved in nitrogen fixation. The C-terminus of the leader peptide includes the prototypical Gly-Gly cleavage motif, where the leader peptide is removed for complete maturation of the peptide product, resulting in an 18 amino acid core with sequence MVTTFFIPSESDDQFFKK. Using the DeepRiPP workflow, EmbA is predicted to be a class II lanthipeptide with 79% class prediction probability, but very low homology to other RiPP cores (Merwin et al., 2020). Threonine and serine residues are present in the core sequence, and could be sites for phosphorylation and dehydration, as is characteristic in lanthipeptide biosynthesis. Nonetheless, cysteines are not found in the core peptide sequence, eliminating the possibility of (methyl)lanthionine macrocycle formation.BLASTp analysis on EmbM shows homology to the dehydration domain of type 2 lanthipeptide biosynthesis proteins, generally termed LanM, which are bifunctional enzymes capable of catalyzing dehydration and cyclization reactions on their substrates (Repka et al., 2017). EmbM was modelled using Phyre2 (Kelley et al., 2015). In accordance with BLAST results, EmbM shows predicted structural similarity to protein kinases and to the dehydratase domain of CylM (83% coverage of the sequence, modelled with 100% confidence), the Enterococcal lanthipeptide synthetase enzyme from the cytolysin biosynthetic pathway (Figure S15A-B) (Dong et al., 2015). EmbM, however, does not contain the typical Zn-dependent cyclization domain which catalyzes the lanthionine macrocycle formation between the thiol side chain of Cys and dehydroamino acids. This observation is consistent with the lack of cysteine residues in the precursor peptide EmbA. Notably, biosynthetic enzymes homologous to the dehydration domain of LanM and lacking the cyclization domain are also present in the biosynthetic clusters for the polytheonamides and aeronamides, highly cytotoxic compounds from '*Candidatus* Entotheonella factor' and *Microvirgula aerodenitrificans* (Bhushan et al., 2019; Freeman et al., 2012, 2016). These enzymes, PoyF and AerF, were shown to dehydrate threonines at core position 1.

In CylM, residues required for phosphorylation are Lys274, Asp347, His349, Asn352, and Asp364 (Dong et al., 2015). In EmbM, sequence alignment and structural modelling allows us to map these residues to Lys178, Asp252, His254, Asn257, and Asp269, suggesting

EmbM's ability to phosphorylate Thr or Ser residues in the precursor peptide core (Figure S15C-D). Mutagenesis investigations in representative lanthipeptide synthetases and CylM have identified residues Lys274, Asp252, His254, Arg506 and Thr512 as key for facilitating phosphate elimination (Dong et al., 2015; Ma et al., 2014; Repka et al., 2017; You and van der Donk, 2007). Lys274, Asp252 and His254 are proposed to both activate the phosphate group for transfer from ATP and stabilize the phosphate in the elimination step. In EmbM, these correspond to Lys178 and Asp155. Notably, EmbM is missing the equivalent of H254, and this residue corresponds to a Pro in EmbM.

Arg506 and Thr512 are proposed to directly assist phosphate elimination in CylM. Thr512 is thought to act as a general base to deprotonate the α-carbon of the phosphorylated residue, generating an enolate, which is stabilized by the side chain of Arg506. In EmbM, CylM's Arg506 is mapped to Arg390 and CylM's Thr512 to Ser396 (Figure S15C-D). Although highly unusual, Thr to Ser mutation is not unprecedented: in *Nostoc sp.* 106C, a LanM enzyme (WP_086758087) with a Ser as predicted general base is associated with a Nif11 precursor peptide (WP_086758085). To our knowledge, the function of this particular LanM enzyme has not been experimentally validated.

Immediately downstream of *embM* is *orf3*. The resulting small protein, which we termed EmbI, shows only distant homology to proteins of unknown function. Notably, EmbI shows no similarity to PqqD, the prototypical RiPP recognition element (RRE) (Burkhart et al., 2015; Kloosterman et al., 2020). A prediction of secondary structural elements of EmbI using PSIPRED and analysis using transmembrane hidden Markov model (TMHHM), indicates the EmbI is comprised of two alpha helices, of which the C-terminal is predicted to be a transmembrane helix (Kandathil et al., 2019; Krogh et al., 2001). Along with the high calculated isoelectric point and proximity to bacteriocin biosynthetic genes, an immunity related function is suggested for EmbI. The scaffold containing *embA* and *embM* also harbors open reading frames likely involved in regulation via a two-component system. Orf1 is predicted to act as a sensory histidine kinase and Orf2 shows high homology to response regulators. The C-terminus of Orf6 shows distant homology to peptidase family C40, often involved in cell wall degradation or remodeling (Aramini et al., 2008). Finally, Orf7 shows homology to glutamine aminotransferase, which takes part in the biosynthesis of guanosine nucleotides. Typical transporters associated with Nif11-type precursor peptides are not present in the cluster neighborhood.

Functional characterization

In order to functionally characterize cluster 75.1, heterologous co-expression experiments and *in vitro* enzymatic assays were conducted. Heterologous co-expression in *E. coli* was initially performed with the constructs pACYCDuet-1-EmbA(MCS1) in *E. coli* BL21(DE3), as a negative control; and pACYCDuet-1-EmbA(MCS1) + pCDFDuet-1-EmbM(MCS2) in *E. coli* BL21(DE3) (Table S5). Expression of both EmbA and EmbM, followed by nickel-affinity chromatography for purification of EmbA resulted in high expression levels for both proteins. Proteolysis with trypsin followed by HPLC-MS analysis resulted in the appearance of two new peaks in the chromatogram upon comparison with an experiment in which only EmbA was expressed. The new peaks displayed monoisotopic masses $(M+3H)^{3+}$ 1182.8769 and $(M+3H)^{3+}$ 1209.5277, which correspond to a difference of +79.9743 and and +159.9267, respectively, in relation to the unmodified trypsin digest fragment of EmbA (Table S5). This mass difference corresponds to the single and double phosphorylation of EmbA (monophosphorylated EmbA calculated mass $(M+3H)^{3+}$ 1182.8685; double phosphorylated EmbA calculated mass $(M+3H)^{3+}$ 1209.5239, Table S5). HPLC-MS/MS was utilized to localize the modifications in the peptide sequence. Fragment ions y16, y15, y14 and b17, b16, b15 allow us to map modifications to Thr(3) and Thr(4) for monophosphorylated products and both Thr(3) and Thr(4) for doubly phosphorylated products (Figure S16A, Table S5). Fragment ions corresponding to Thr(4) phosphorylation were more abundant, pointing to initial modification at that position being favored. Cleavage of EmbA with LahT150 (Bobeica et al., 2019) resulted in the removal of the leader peptide and identification of mono and double phosphorylated peptide cores, in line with the results obtained with trypsin (Table S5).

*In silico* analysis of EmbM suggests the enzyme is capable of phosphorylation of substrate peptides, given all residues proposed to be involved in kinase activity of CylM can be mapped to EmbM. This hypothesis is confirmed experimentally with the results described above. Given the broad similarities of EmbM to AerF, PoyF and the dehydration domain of LanM enzymes, β-elimination of phosphate groups on Thr(3) and Thr(4) to yield dehydrated products is also expected. Only trace amounts of dehydrated EmbA were found in the initial experiments described above. A number of additional co-expression conditions were tested: *E. coli* expression hosts were varied, along with expression media, temperature and time (Table S5). Different construct combinations were also tested: precursor and EmbM were expressed in their genetic context or expressed from individual plasmids with different copy numbers and the genes encoding each protein were also codon-optimized for *E. coli* expression. EmbI was also included in co-expression experiments, given its putative role in providing the native producing organism, and likely the heterologous expression host, with immunity to the biological activity of the cluster product. All conditions assayed resulted in the same outcome: high amounts of phosphorylated products were detected, but dehydration was not observed.

Co-expression of EmbA and EmbM was also performed in *Microvirgula aerodenitrificans*, a Betaproteobacterium isolated from activated sludge (Bhushan et al., 2019). We hypothesized that, due to the similarities between EmbM and AerF and the fact that *M. aerodenitrificans* is also found in aquatic environments, EmbM was likely to display its full predicted activity. Nonetheless, 94% of the precursor peptide was converted to phosphorylated products (percent conversion was calculated based on the relative peak areas in extracted ion chromatograms). *In vitro* enzymatic assays with EmbA and EmbM were also conducted. The proteins were individually purified and assayed with $MgCl_2$ and ATP as a co-substrate. A variety of conditions were screened and, upon incubation of EmbA and EmbM at 37°C for 3 days, upwards of 75% conversion to phosphorylated products was observed, in line with the results of co-expression experiments.

Intriguingly, efficient phosphorylation by wild-type LanM-type enzymes without concomitant β-elimination and generation of dehydrated amino acids is, to our knowledge, unprecedented. Two hypotheses can be proposed to explain this outcome. First, metagenomic DNA from the producing organism *Ca*. E. malaspinii was sampled from bathypelagic zones, where extreme pressure, low temperatures and high salinity likely have an effect on metabolic processes and adaptation. Therefore, it is possible that the conditions required for complete predicted activity of EmbM cannot be reproduced in the laboratory. Second, EmbM is missing one of the three residues thought to stabilize the phosphate group for elimination and, notably, the predicted base that deprotonates the α-carbon of phosphorylated residues is a Ser as opposed to a more common Thr. On principle, however, these amino acid modifications are unlikely to have such a drastic effect on enzyme activity.

The biosynthesis of the hypothetical dehydrated final product of cluster 75.1 was recreated *in vitro* by combining enzymatic and chemical methods. Single and double phosphorylated products were generated by reaction of EmbA and EmbM. Cleavage of the leader peptide with the promiscuous LahT150 protease yielded the phosphorylated core peptide. Finally, β-elimination of phosphorylated peptides was achieved by treatment with base. Production of single and double dehydrated products was confirmed by HR-MS and HR-MS/MS (Table S5, Figure S16B). Further confirmation of the presence of α,β-unsaturated Thr residues resulted from derivatization of EmbA trypsin fragments with DTT (Table S5, Figure S16B). Dehydrobutyrine residues are more stable in the *Z* configuration and, with the exception of cypemycin, all Dhb-containing RiPPs harbor the *Z* isomer (Butler et al., 2018; Dugave and Demange, 2003; Siodłak, 2015). We thus depict Dhb(3) and Dhb(4) in the *Z* configuration.

The presence of dehydrated residues in peptides impart significant structural changes and lead to interesting biological activities. Linear dehydrated peptides such as cypemycin (Komiyama et al., 1993), salinipeptins (Shang et al., 2019), albopeptide (Wang et al., 2020), loihichelins (Homann et al., 2009), bogorol (Barsby et al., 2001) and lavendomycin (Komori et al., 1985) show activities mostly as antibiotics, and, more rarely, as siderophores or anti-cancer compounds. This literature precedent, together with the narrow-spectrum antibiotic activity typical of bacteriocins and the putative immunity function of EmbI, allow us to hypothesize the ecological function of 75.1 as a growth inhibitor of strains related to the producing deep sea *Ca*. Eudoremicrobiaceae (*Ca*. E. malaspinii). Bacteria related to *Ca*. E. malaspinii are as-yet uncultured including the three closest related phyla: *Armatimonadata*, UBP13, and UBP9 (Figure S4). Therefore, we were only able to test the biological activity of the dehydrated peptide against eight common laboratory strains of bacteria and fungi as well as four strains isolated from marine environments (Table S5). No growth inhibition was observed for the strains tested. We also tested the antibiotic activity of phosphorylated peptides against *E. coli, S. aureus, A. baumanii, and P. aeruginosa* and, similarly, no growth inhibition was observed.

### Supplementary Figures

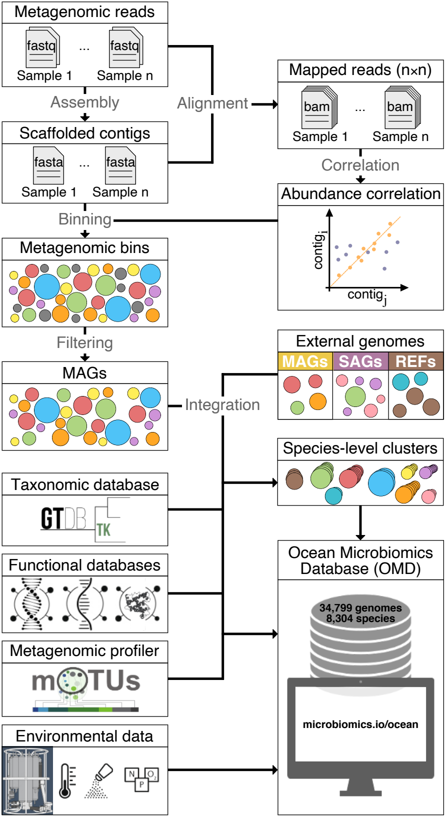

Metagenomic reads
fastq
...
fastq
Sample 1
Sample n
Assembly
Alignment
Mapped reads (n×n)
bam
...
bam
Sample 1
Sample n
Scaffolded contigs
fasta
...
fasta
Sample 1
Sample n
Correlation
Abundance correlation
Binning
contig_i
contig_j
Metagenomic bins
Filtering
MAGs
Integration
External genomes
MAGs
SAGs
REFs
Species-level clusters
Taxonomic database
GTDB TK
Functional databases
Ocean Microbiomics Database (OMD)
34,799 genomes
8,304 species
Metagenomic profiler
mOTUs
microbiomics.io/ocean
Environmental data
N
P
O2

**Figure S1: Overview of the bioinformatic pipeline. Related to Figure 1.**

Quality-controlled, high-throughput DNA sequencing reads from ocean microbial community samples were individually assembled into metagenomic scaffolded contigs (scaffolds). Sequencing reads from large subsets (n ranging from 58 to 610) of all samples were aligned to scaffolds of each individual sample to compute relative copy-number abundances for each scaffold in each sample. Based on a combination of tetranucleotide frequency, within-sample co-abundance and between-sample abundance correlations, scaffolds were grouped into a total of 62,874 metagenomic bins, each with total nucleotide sequence lengths of >200 kbp. These metagenomic bins were filtered for genome completeness and contamination, resulting in 26,293 metagenome assembled genomes (MAGs). These MAGs were complemented with external sets of MAGs, single amplified genomes (SAGs) and genomes from cultured isolates (REFs). The combined set of 34,799 genomes was clustered at the species level using a 95% average nucleotide identity (ANI) and, along with taxonomic and functional annotations, abundance profiles and contextual information, compiled into the Ocean Microbiomics Database (OMD); see methods for details (Methods).

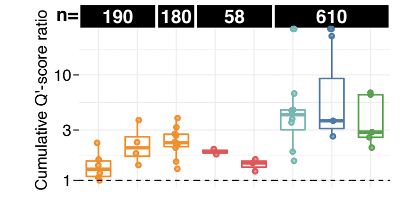

n=
190
180
58
610
Cumulative Q'-score ratio
10
3
1

**Figure S2: Impact of abundance correlation on MAGs recovery and quality. Related to Figure 1.**

In this study, MAGs were reconstructed using abundance correlation information (Figure S1) (Methods), which resulted in both higher cumulative quality scores per sample and individual quality scores per MAG. The ratio of cumulative quality scores (Supplemental information) of MAGs binned with and without differential coverage information was on average (median) 2.3 across the different datasets. Per individual MAGs, a mean quality score increase of 20% was achieved. The number of samples used for differential coverage profiling are indicated above the boxplots. The colors of the boxplots reflect the different datasets as indicated in Figure 1C.

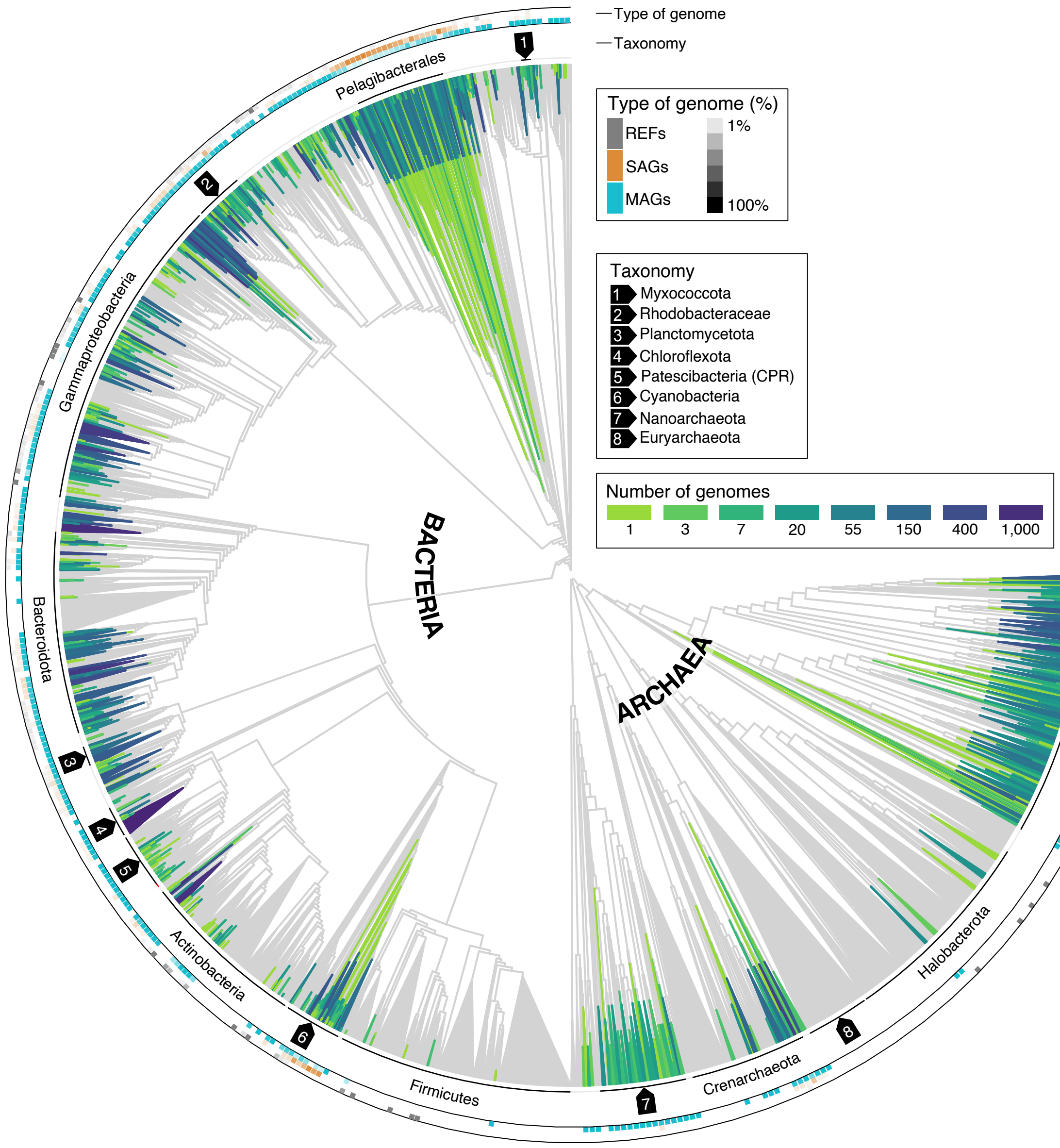

Type of genome
Taxonomy
Type of genome (%)
REFs
SAGs
MAGs
1%
100%
Taxonomy
1 Myxococcota
2 Rhodobacteraceae
3 Planctomycetota
4 Chloroflexota
5 Patescibacteria (CPR)
6 Cyanobacteria
7 Nanoarchaeota
8 Euryarchaeota
Number of genomes
1
3
7
20
55
150
400
1,000
BACTERIA
ARCHAEA
Pelagibacterales
Gammaproteobacteria
Bacteroidota
Actinobacteria
Firmicutes
Crenarchaeota
Halobacterota
Poseidoniia

**Figure S3: Different genome reconstruction strategies capture complementary phylogenomic diversity. Related to figure 1.**

Reconstructed MAGs, external MAGs, SAGs as well as REFs detected across the set of 1,038 ocean metagenomes were placed on the GTDB backbone trees (Parks et al., 2018) revealing that the different genome types (MAGs, SAGs and REFs) capture complementary phylogenomic diversity. Similar to Figure 3, the green-to-blue colors of the branches indicate the number of genomes in that part of the tree. The inner layer denotes the taxonomy of specific clades (some indicated by arrows due to limited space). The outer layer represents the percentage of genomes across the binned tree for each genome type. Clades without any genome from the OMD were left in gray. For visualization purposes, the last 15% of the nodes are collapsed.

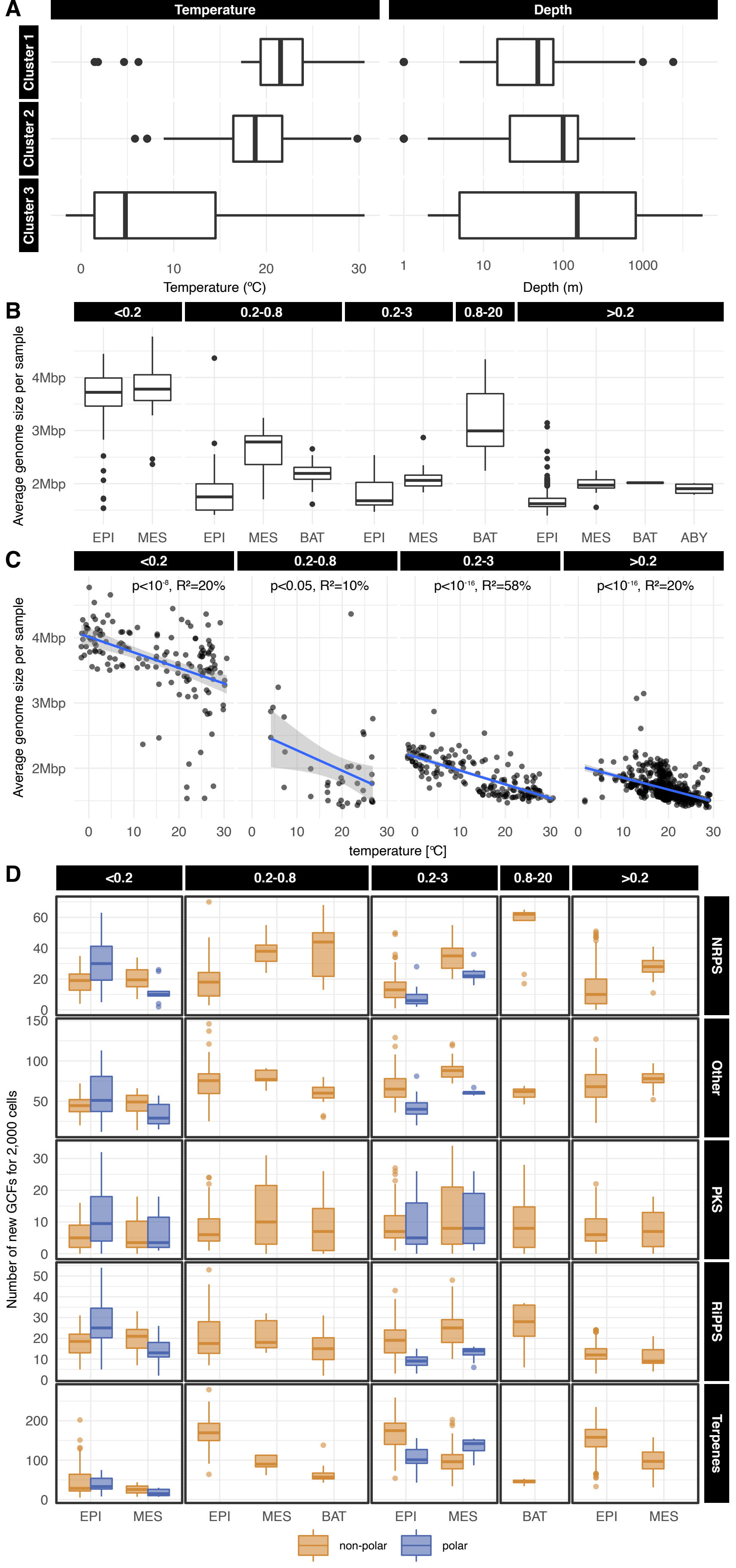
A
Temperature
Depth
Cluster 1
Cluster 2
Cluster 3
Temperature (ºC)
Depth (m)
B
<0.2
0.2-0.8
0.2-3
0.8-20
>0.2
Average genome size per sample
EPI
MES
BAT
ABY
C
p<10⁻⁸, R²=20%
p<0.05, R²=10%
p<10⁻¹⁶, R²=58%
p<10⁻¹⁶, R²=20%
temperature [ºC]
D
Number of new GCFs for 2,000 cells
NRPS
Other
PKS
RiPPS
Terpenes
non-polar
polar

**Figure S4: Environmental drivers and novelty of the ocean microbiome biosynthetic potential. Related to Figure 2.**

(A) We found temperature and depth to be significantly different between the sample clusters identified based on biosynthetic potential composition (FDR corrected pairwise Wilcoxon tests) (Figure 2C). (B, C) The average genome size per sample was significantly larger in deeper waters (FDR corrected pairwise Wilcoxon tests) and was inversely correlated with temperature (linear model). (D) We estimated the discovery potential of different microbial communities by counting the number of new GCFs (Methods) detected in a sample after rarefaction of per-cell GCFs abundance profile to 2,000 cells. Although well studied communities (non-polar epipelagic prokaryote-enriched (0.2-3 μm) and virus-depleted (>0.2 μm)) displayed the highest discovery potential for terpenes, least explored communities (polar, deep, virus- and particle-enriched) were found to have the highest potential for NRPS, PKS, RiPPs or other natural products discovery. Polar is defined as absolute latitude >60º. NRPS: Non-Ribosomal Peptide Synthetases; PKS: Polyketide Synthase; RiPP: Ribosomally Synthesized and Post-translationally modified Peptide.

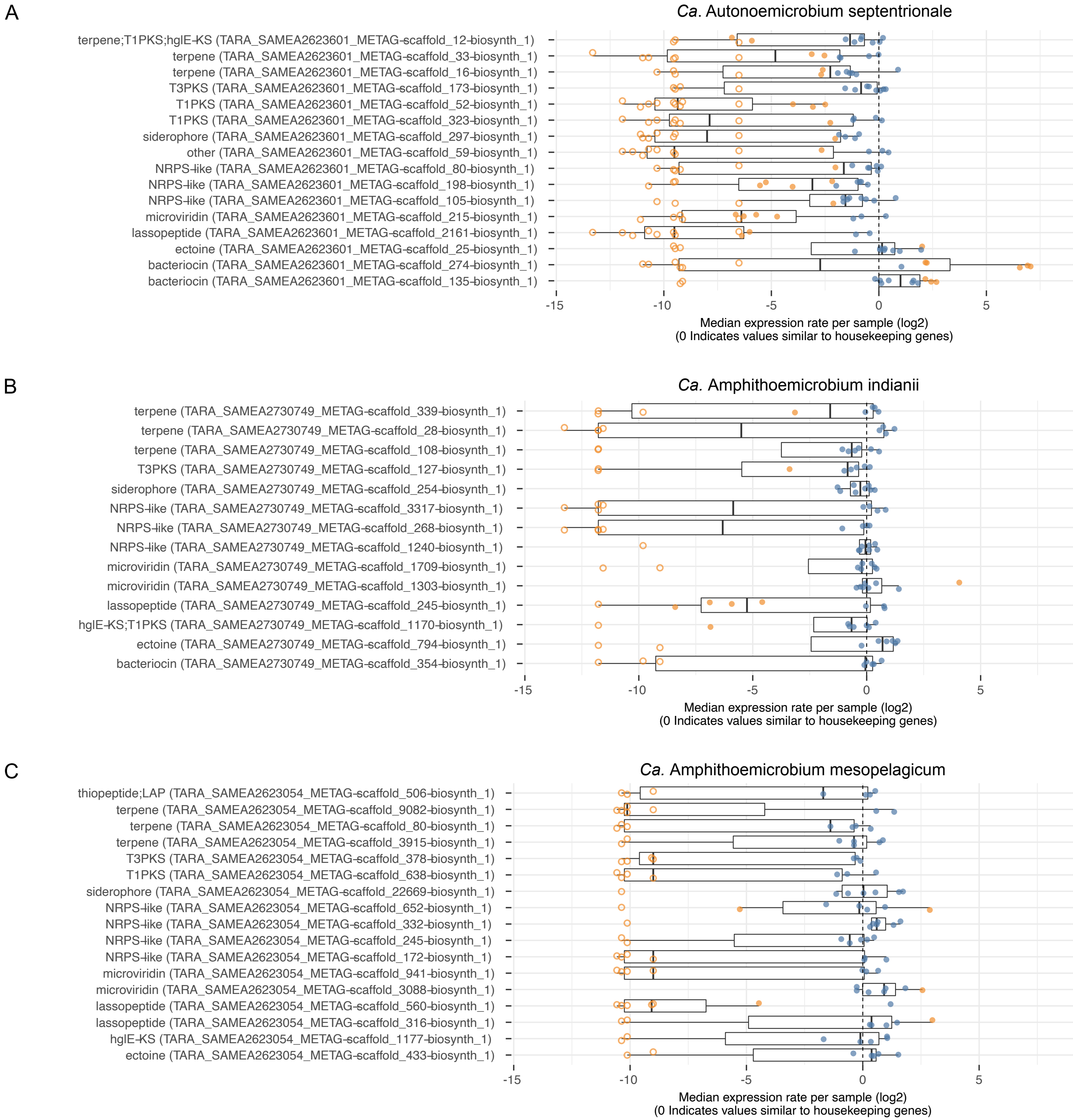
A
Ca. Autonoemicrobium septentrionale
terpene;T1PKS;hglE-KS (TARA_SAMEA2623601_METAG-scaffold_12-biosynth_1)
terpene (TARA_SAMEA2623601_METAG-scaffold_33-biosynth_1)
terpene (TARA_SAMEA2623601_METAG-scaffold_16-biosynth_1)
T3PKS (TARA_SAMEA2623601_METAG-scaffold_173-biosynth_1)
T1PKS (TARA_SAMEA2623601_METAG-scaffold_52-biosynth_1)
T1PKS (TARA_SAMEA2623601_METAG-scaffold_323-biosynth_1)
siderophore (TARA_SAMEA2623601_METAG-scaffold_297-biosynth_1)
other (TARA_SAMEA2623601_METAG-scaffold_59-biosynth_1)
NRPS-like (TARA_SAMEA2623601_METAG-scaffold_80-biosynth_1)
NRPS-like (TARA_SAMEA2623601_METAG-scaffold_198-biosynth_1)
NRPS-like (TARA_SAMEA2623601_METAG-scaffold_105-biosynth_1)
microviridin (TARA_SAMEA2623601_METAG-scaffold_215-biosynth_1)
lassopeptide (TARA_SAMEA2623601_METAG-scaffold_2161-biosynth_1)
ectoine (TARA_SAMEA2623601_METAG-scaffold_25-biosynth_1)
bacteriocin (TARA_SAMEA2623601_METAG-scaffold_274-biosynth_1)
bacteriocin (TARA_SAMEA2623601_METAG-scaffold_135-biosynth_1)
-15
-10
-5
0
5
Median expression rate per sample (log2)
(0 Indicates values similar to housekeeping genes)
B
Ca. Amphithoemicrobium indianii
terpene (TARA_SAMEA2730749_METAG-scaffold_339-biosynth_1)
terpene (TARA_SAMEA2730749_METAG-scaffold_28-biosynth_1)
terpene (TARA_SAMEA2730749_METAG-scaffold_108-biosynth_1)
T3PKS (TARA_SAMEA2730749_METAG-scaffold_127-biosynth_1)
siderophore (TARA_SAMEA2730749_METAG-scaffold_254-biosynth_1)
NRPS-like (TARA_SAMEA2730749_METAG-scaffold_3317-biosynth_1)
NRPS-like (TARA_SAMEA2730749_METAG-scaffold_268-biosynth_1)
NRPS-like (TARA_SAMEA2730749_METAG-scaffold_1240-biosynth_1)
microviridin (TARA_SAMEA2730749_METAG-scaffold_1709-biosynth_1)
microviridin (TARA_SAMEA2730749_METAG-scaffold_1303-biosynth_1)
lassopeptide (TARA_SAMEA2730749_METAG-scaffold_245-biosynth_1)
hglE-KS;T1PKS (TARA_SAMEA2730749_METAG-scaffold_1170-biosynth_1)
ectoine (TARA_SAMEA2730749_METAG-scaffold_794-biosynth_1)
bacteriocin (TARA_SAMEA2730749_METAG-scaffold_354-biosynth_1)
-15
-10
-5
0
5
Median expression rate per sample (log2)
(0 Indicates values similar to housekeeping genes)
C
Ca. Amphithoemicrobium mesopelagicum
thiopeptide;LAP (TARA_SAMEA2623054_METAG-scaffold_506-biosynth_1)
terpene (TARA_SAMEA2623054_METAG-scaffold_9082-biosynth_1)
terpene (TARA_SAMEA2623054_METAG-scaffold_80-biosynth_1)
terpene (TARA_SAMEA2623054_METAG-scaffold_3915-biosynth_1)
T3PKS (TARA_SAMEA2623054_METAG-scaffold_378-biosynth_1)
T1PKS (TARA_SAMEA2623054_METAG-scaffold_638-biosynth_1)
siderophore (TARA_SAMEA2623054_METAG-scaffold_22669-biosynth_1)
NRPS-like (TARA_SAMEA2623054_METAG-scaffold_652-biosynth_1)
NRPS-like (TARA_SAMEA2623054_METAG-scaffold_332-biosynth_1)
NRPS-like (TARA_SAMEA2623054_METAG-scaffold_245-biosynth_1)
NRPS-like (TARA_SAMEA2623054_METAG-scaffold_172-biosynth_1)
microviridin (TARA_SAMEA2623054_METAG-scaffold_941-biosynth_1)
microviridin (TARA_SAMEA2623054_METAG-scaffold_3088-biosynth_1)
lassopeptide (TARA_SAMEA2623054_METAG-scaffold_560-biosynth_1)
lassopeptide (TARA_SAMEA2623054_METAG-scaffold_316-biosynth_1)
hglE-KS (TARA_SAMEA2623054_METAG-scaffold_1177-biosynth_1)
ectoine (TARA_SAMEA2623054_METAG-scaffold_433-biosynth_1)
-15
-10
-5
0
5
Median expression rate per sample (log2)
(0 Indicates values similar to housekeeping genes)

**Figure S5: Expression of *Ca*. Eudoremicrobiaceae BGCs. Related to Figure 4.**

All BGCs encoded by *Ca*. Autonoemicrobium septentrionale (A), *Ca*. Amphithoemicrobium indianii (B) and *Ca*. Amphithoemicrobium mesopelagicum representatives were found to be expressed in the natural environment (in the 623 *Tara* Oceans metatranscriptomic samples (Salazar et al. 2019; Carradec et al. 2018; Methods). Some displayed near constitutive expression while others appear to be tightly regulated across the metatranscriptomes studied here. Filled circles indicate samples where active transcription was detected. Orange data points indicate values below or above a $\log_2$ fold change from the constitutive expression rate of housekeeping genes. All the BGCs encoded by *Ca*. E. taraoceanii were also found to be expressed (Figure 4C). The expression of *Ca*. E. malaspinii BGCs could not be investigated since that species was not sufficiently abundant in the epipelagic and mesopelagic ocean, the only layers for which metatranscriptomes were available.

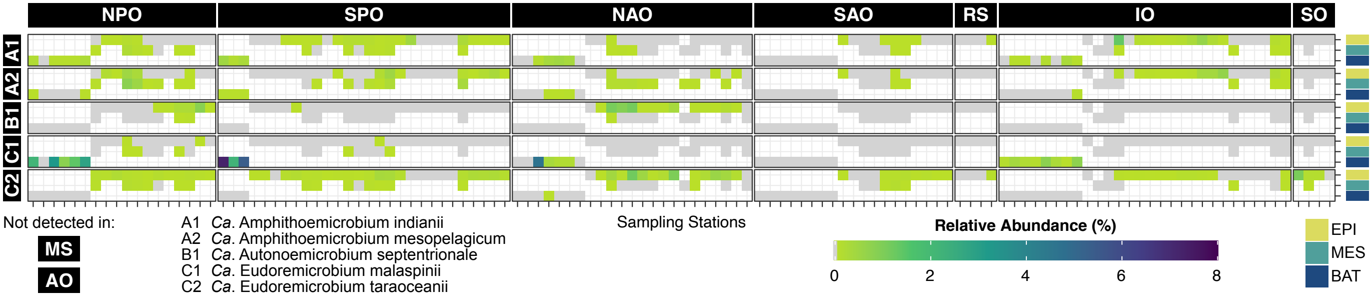
NPO
SPO
NAO
SAO
RS
IO
SO
A1
A2
B1
C1
C2
Sampling Stations
Not detected in:
MS
AO
A1 Ca. Amphithoemicrobium indianii
A2 Ca. Amphithoemicrobium mesopelagicum
B1 Ca. Autonoemicrobium septentrionale
C1 Ca. Eudoremicrobium malaspinii
C2 Ca. Eudoremicrobium taraoceanii
Relative Abundance (%)
0
2
4
6
8
EPI
MES
BAT

**<u>Figure S6: Ecological distribution of *Ca*. Eudoremicrobiaceae spp. in the ocean. Related to Figure 4.</u>**

Relative abundance (profiled using the extended mOTUs database, see Methods) profiles showed *Ca*. Eudoremicrobiaceae spp. to be abundant and prevalent in most ocean basins as well as throughout the water column (from the surface down to at least 4,000 meters deep). Based on these estimations, we found *Ca*. E. malaspinii to make up to 7% of the prokaryotic cells in bathypelagic particle-associated communities. We considered a species present at a station if the mOTUs cluster was detected in any of the size fraction of a given depth layer. NPO: North Pacific Ocean; SPO: South Pacific Ocean; NAO: North Atlantic Ocean; SAO: South Atlantic Ocean; RS: Red Sea; IO: Indian Ocean; SO: Southern Ocean; MS: Mediterranean Sea; AO: Arctic Ocean.

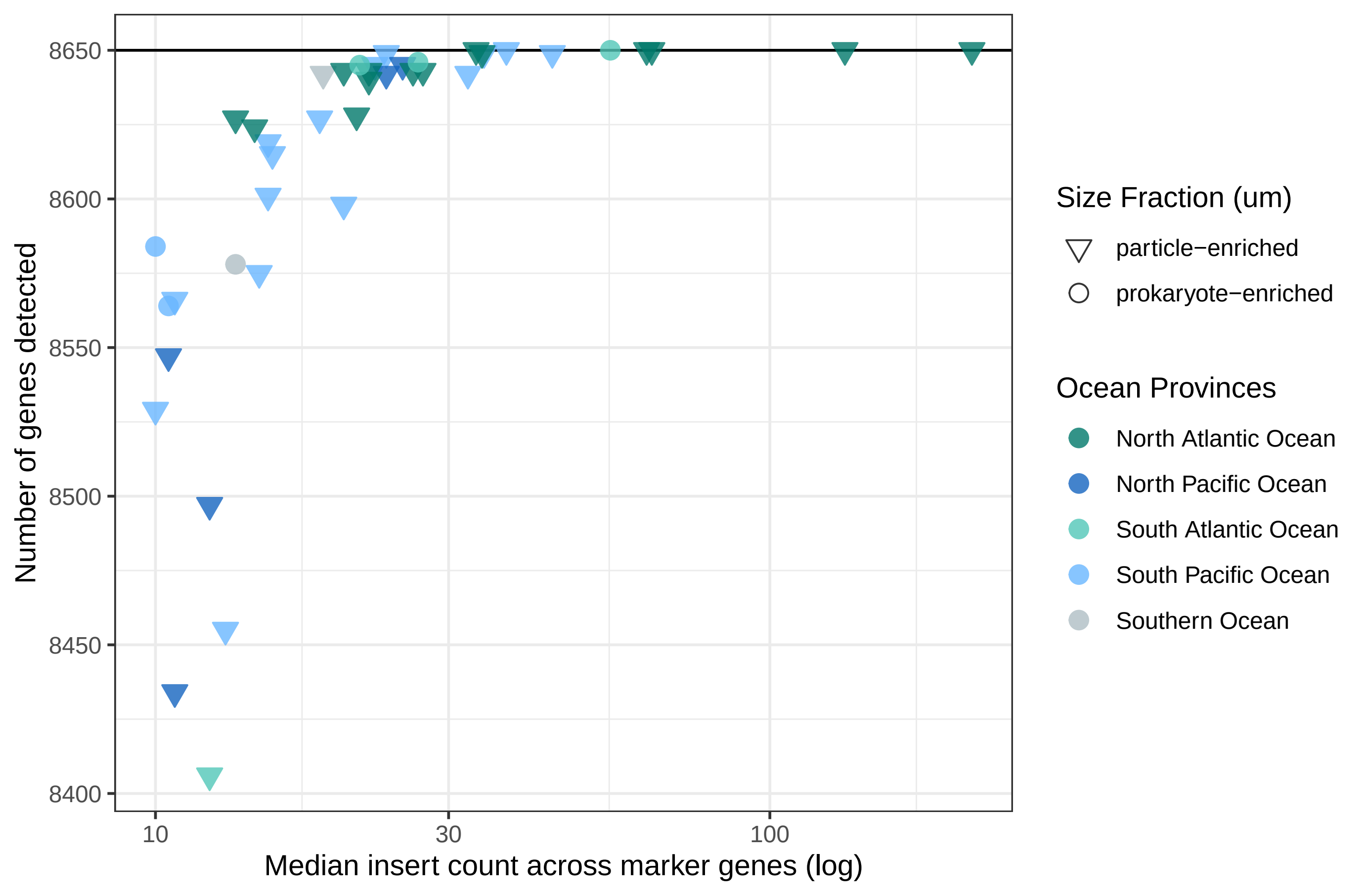
Number of genes detected
8650
8600
8550
8500
8450
8400
10
30
100
Median insert count across marker genes (log)
Size Fraction (um)
particle−enriched
prokaryote−enriched
Ocean Provinces
North Atlantic Ocean
North Pacific Ocean
South Atlantic Ocean
South Pacific Ocean
Southern Ocean

**Figure S7: Stable gene content in *Ca*. E. taraoceanii natural populations. Related to Figure 4.**

We investigated the metagenomic detection of the 8,500 genes encoded by the *Ca*. E. taraoceanii representative, using methodology identical to the transcriptomic analyses (Methods). In samples where the 10 marker genes were detected, we counted the number of genes with one or more insert(s). We found that the 8,500 genes were detected in several ocean basins and different size fractions, with variation in detection rates likely due to variable sequencing depths across samples and datasets. This indicates, at least for the gene set covered by the reconstructed genome, that niche partitioning may be driven by gene expression changes rather than gene content variation.

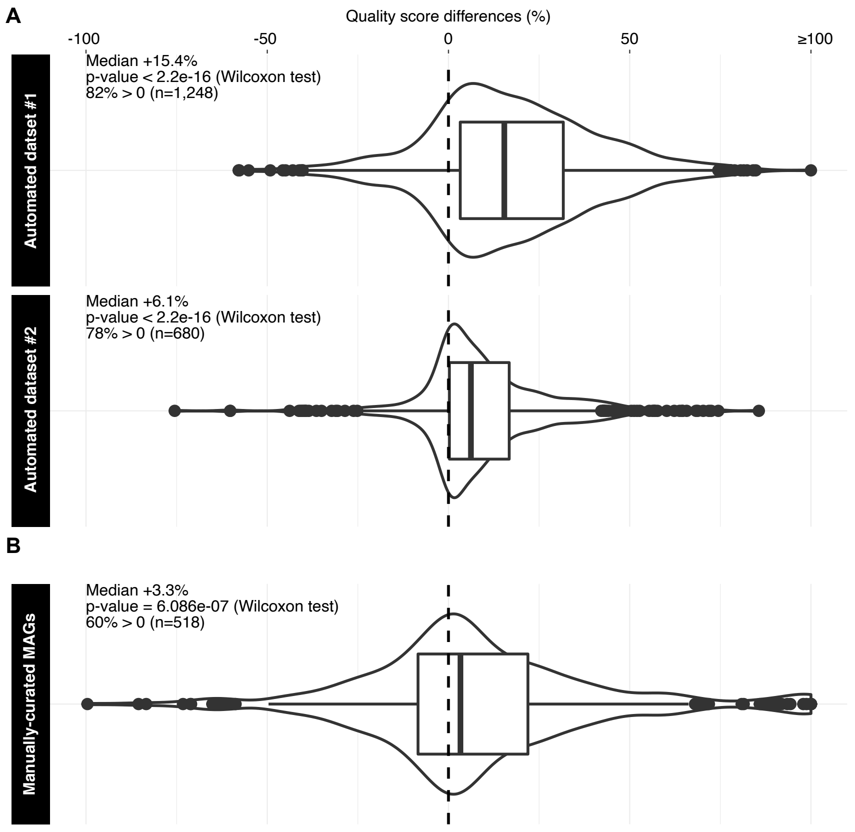

A
Quality score differences (%)
-100
-50
0
50
≥100
Automated dataset #1
Median +15.4%
p-value < 2.2e-16 (Wilcoxon test)
82% > 0 (n=1,248)
Automated dataset #2
Median +6.1%
p-value < 2.2e-16 (Wilcoxon test)
78% > 0 (n=680)
B
Manually-curated MAGs
Median +3.3%
p-value = 6.086e-07 (Wilcoxon test)
60% > 0 (n=518)

**Figure S8: Quality improvement over other ocean MAGs datasets. Related to Figure 1.**

By benchmarking the quality of the MAGs reconstructed in this study (Supplemental information), we found that combining single-sample assemblies with large-scale abundance correlations achieved on average significantly higher community-defined quality scores (Bowers et al., 2017) than and (A) two datasets of automatically generated MAGs, dataset #1 (Tully et al., 2018) and dataset #2 (Parks et al., 2017), and (B) even manually curated MAGs (Delmont et al., 2018). ‘n’ denotes the number of possible comparisons (i.e. number of shared species) with the different MAGs sets.

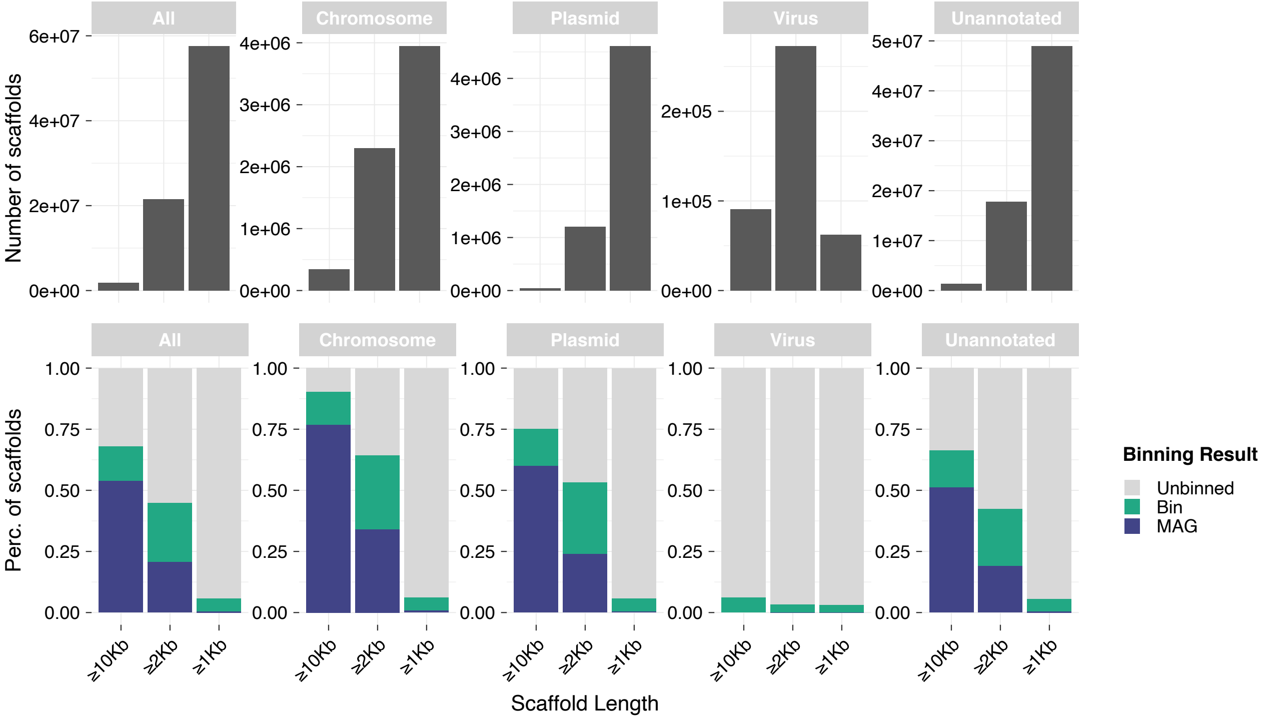

All
Chromosome
Plasmid
Virus
Unannotated
Number of scaffolds
0e+00
2e+07
4e+07
6e+07
1e+06
2e+06
3e+06
4e+06
1e+05
2e+05
1e+07
3e+07
5e+07
Perc. of scaffolds
0.00
0.25
0.50
0.75
1.00
≥10Kb
≥2Kb
≥1Kb
Scaffold Length
Binning Result
Unbinned
Bin
MAG

**Figure S9: Binning proportion of different metagenomic fragments. Related to Figure 1.**

We investigated the bin membership of >80 M scaffolds across size and fragment type. These scaffolds were annotated to identify chromosomes, plasmids and phages (Supplemental information). The difference between chromosomes and plasmids binning rates provides an evaluation of the bias of the MAG reconstruction against hypervariable regions within the genomes. Annotations were integrated to classify scaffolds as follows, **chromosomes** (‘*eukrep = Prokarya & plasflow prediction = chromosome & cbar prediction = Chromosome & plasmidfinder plasmid = NaN & deepvirfinder pvalue > 0.05 & virsorter score = NaN*’), **plasmids** (‘*(plasmidfinder plasmid != NaN* | *(plasflow prediction = plasmid & cbar prediction = Plasmid)) & eukrep = Prokarya & virsorter score not in [1, 2] & deepvirfinder pvalue > 0.05*’), **viruses** (‘*virsorter score >= 1 & deepvirfinder pvalue < 0.01 & eukrep = Prokarya & plasflow prediction != plasmid & cbar prediction != Plasmid*’) or **unannotated**.

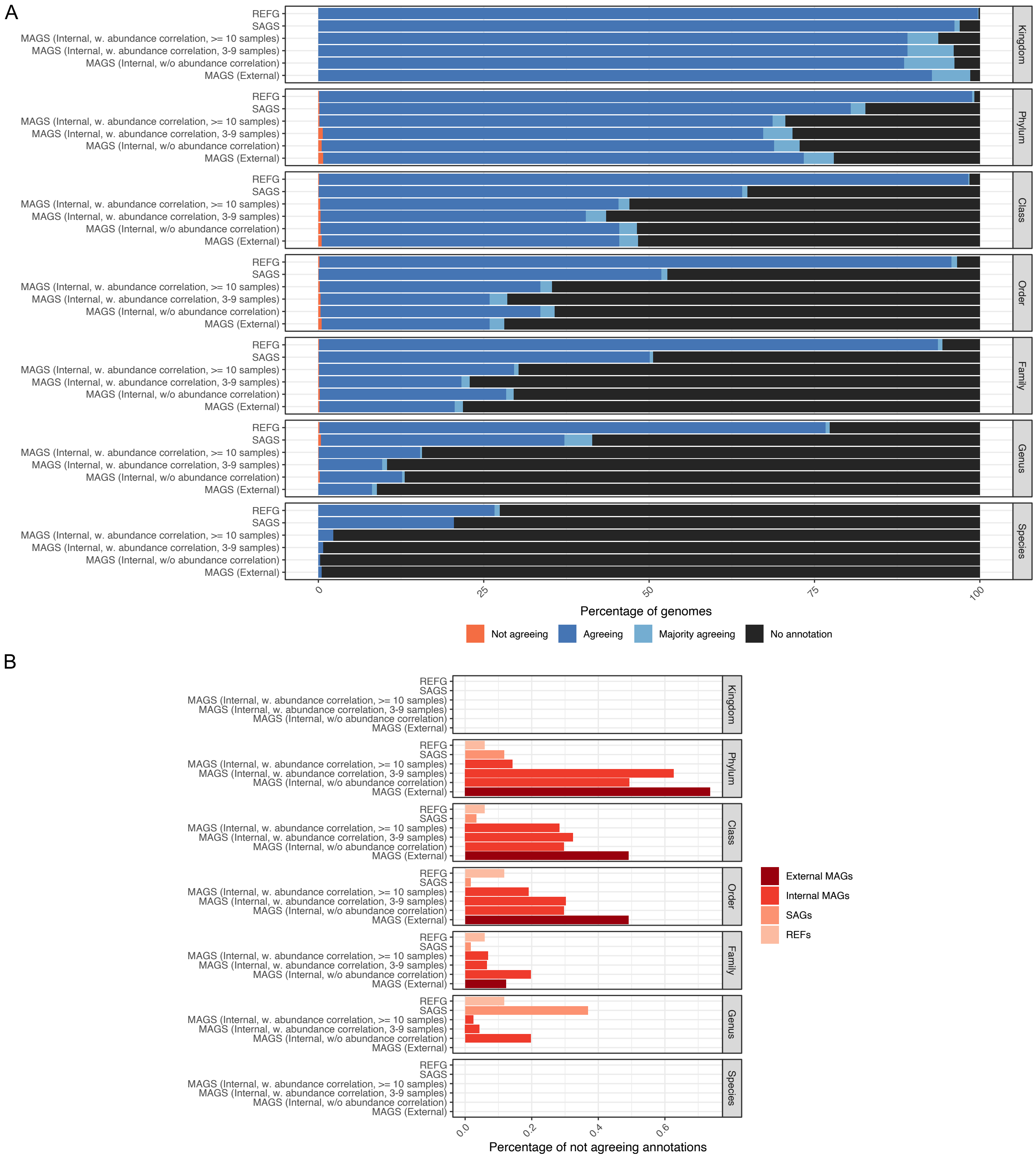

A
REFG
SAGS
MAGS (Internal, w. abundance correlation, >= 10 samples)
MAGS (Internal, w. abundance correlation, 3-9 samples)
MAGS (Internal, w/o abundance correlation)
MAGS (External)
Kingdom
Phylum
Class
Order
Family
Genus
Species
0
25
50
75
100
Percentage of genomes
Not agreeing
Agreeing
Majority agreeing
No annotation
B
0.0
0.2
0.4
0.6
Percentage of not agreeing annotations
External MAGs
Internal MAGs
SAGs
REFs

**Figure S10: Evaluation of genome chimerism. Related to Figure 1.**

All genomes in the extended OMD were evaluated for chimerism using the taxonomic annotation of 10 universal single copy marker genes (Supplemental information). (A) For each taxonomic level, the genomes were classified as: "No annotation" if a maximum of one gene out of 10 was annotated; “Agreeing" if all genes had the same annotation; "Majority agreeing" if more than half agreed and “Not agreeing” otherwise. The evaluation was split for the genomes origin (y-axis). (B) Percentage of “Not agreeing” annotations over all the annotated clades (i.e. the sum of “Agreeing”, Majority agreeing” and “Not agreeing”). Notably, across all MAGs the rate of disagreement was <1% with that rate being ~0.1% for MAGs with differential coverage index ≥10 (i.e. 75% of the MAGs), suggesting the added value of abundance correlation in reducing the rates of chimera.

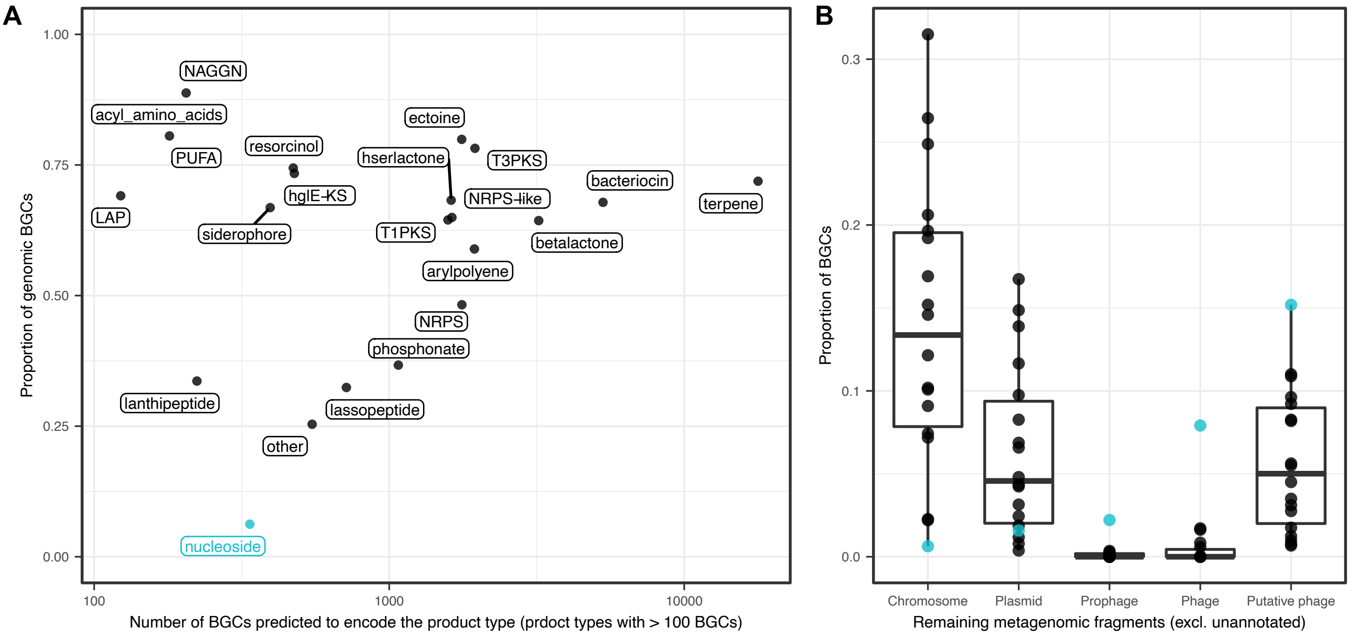
A
NAGGN
acyl_amino_acids
PUFA
resorcinol
hglE-KS
LAP
siderophore
ectoine
hserlactone
T3PKS
NRPS-like
T1PKS
arylpolyene
betalactone
bacteriocin
terpene
NRPS
phosphonate
lanthipeptide
lassopeptide
other
nucleoside
1.00
0.75
0.50
0.25
0.00
Proportion of genomic BGCs
100
1000
10000
Number of BGCs predicted to encode the product type (prdoct types with > 100 BGCs)
B
0.3
0.2
0.1
0.0
Proportion of BGCs
Chromosome
Plasmid
Prophage
Phage
Putative phage
Remaining metagenomic fragments (excl. unannotated)

**<u>Figure S11: Distribution of nucleoside BGCs across genomic and metagenomic fragments. Related to Figure 2.</u>**

(A) Investigation of the proportion of BGCs binned within a MAG by product type showed that nucleosides were most rarely encoded in MAGs. (B) Breakdown by fragment type of the BGCs in the remaining metagenomic fragments. Strikingly, nucleoside BGCs were rarely encoded on predicted chromosome fragments and most often in predicted phage fragments (Supplemental information). For this analysis, we refined the prediction described in Figure S9 with **prophages** (‘*virsorter category = prophage & virsorter score >= 1 & eukrep = Prokarya & plasflow prediction != plasmid & cbar prediction != Plasmid’*), **phages** (‘*virsorter category = phage & virsorter score >= 1 & deepvirfinder pvalue < 0.01 & eukrep = Prokarya & plasflow prediction != plasmid & cbar prediction != Plasmid’*) and **putative phages** (*not in **phages** & ((‘virsorter category = phage & virsorter score >= 1) | deepvirfinder pvalue < 0.05) & eukrep != Eukarya & plasflow prediction != plasmid & cbar prediction != Plasmid’*).

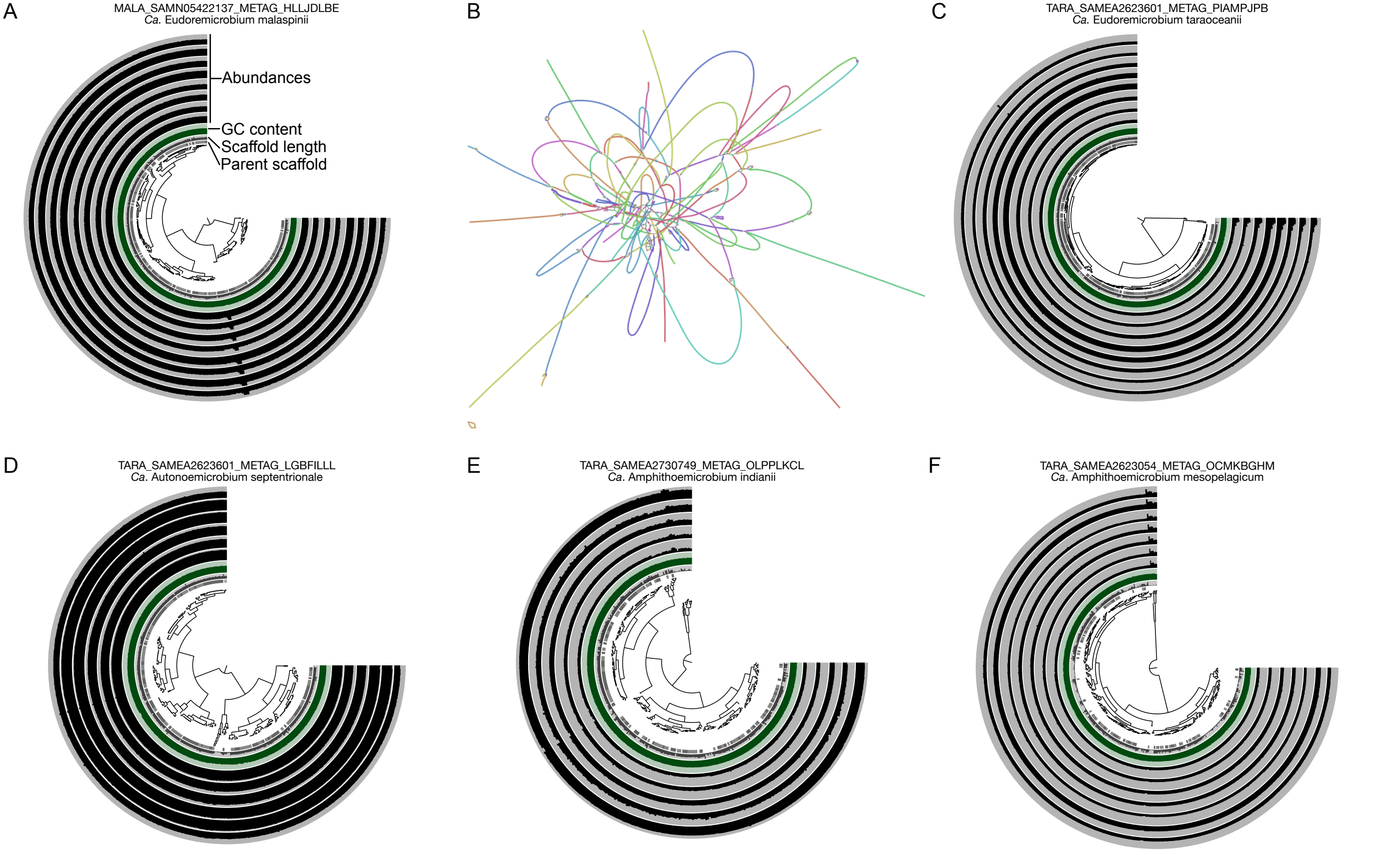

A
MALA_SAMN05422137_METAG_HLLJDLBE
*Ca.* Eudoremicrobium malaspinii
Abundances
GC content
Scaffold length
Parent scaffold
B
C
TARA_SAMEA2623601_METAG_PIAMPJPB
*Ca.* Eudoremicrobium taraoceanii
D
TARA_SAMEA2623601_METAG_LGBFILLL
*Ca.* Autonoemicrobium septentrionale
E
TARA_SAMEA2730749_METAG_OLPPLKCL
*Ca.* Amphithoemicrobium indianii
F
TARA_SAMEA2623054_METAG_OCMKBGHM
*Ca.* Amphithoemicrobium mesopelagicum

**<u>Figure S12: Manual inspection of *Ca.* Eudoremicrobiaceae MAGs. Related to Figure 4.</u>**

(A, C, D, E and F) Anvi'o interface of representatives of the five *Ca*. Eudoremicrobiaceae species reveals stable abundance correlation patterns across the vast majority of the genomes, indicative of low contamination rates (Supplemental information). (B) Inspection of the assembly graph for *Ca*. E. malaspinii (Supplemental information) showed that all scaffolds from the representative genomes were connected with the exception of a single 20 kbp one.

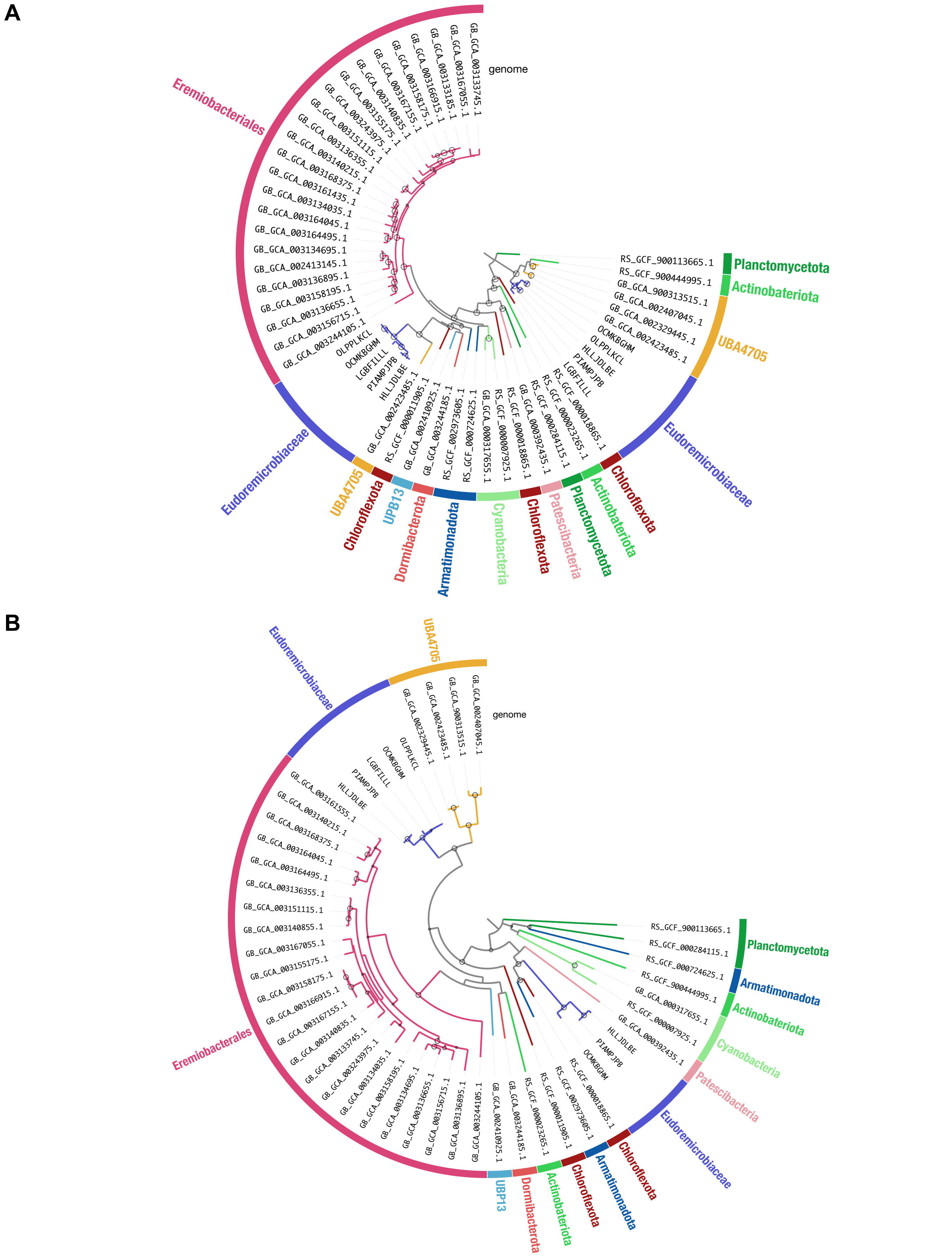
A
genome
GB_GCA_003133745.1
GB_GCA_003167055.1
GB_GCA_003133185.1
GB_GCA_003166915.1
GB_GCA_003158175.1
GB_GCA_003167155.1
GB_GCA_003140835.1
GB_GCA_003155175.1
GB_GCA_003243975.1
GB_GCA_003151115.1
GB_GCA_003136355.1
GB_GCA_003140215.1
GB_GCA_003168375.1
GB_GCA_003161435.1
GB_GCA_003134035.1
GB_GCA_003164045.1
GB_GCA_003164495.1
GB_GCA_003134695.1
GB_GCA_002413145.1
GB_GCA_003136895.1
GB_GCA_003158195.1
GB_GCA_003136655.1
GB_GCA_003156715.1
GB_GCA_003244105.1
OLPPLKCL
OCMKBGHM
LGBFILLL
PIAMPJPB
HLLJDLBE
GB_GCA_002423485.1
RS_GCF_000011905.1
GB_GCA_002410925.1
GB_GCA_003244185.1
RS_GCF_002973605.1
RS_GCF_000724625.1
GB_GCA_000317655.1
RS_GCF_000007925.1
RS_GCF_000018865.1
GB_GCA_000392435.1
RS_GCF_000284115.1
RS_GCF_000023265.1
RS_GCF_000018865.1
LGBFILLL
PIAMPJPB
HLLJDLBE
OLPPLKCL
OCMKBGHM
GB_GCA_002423485.1
GB_GCA_002329445.1
GB_GCA_002407045.1
GB_GCA_900313515.1
RS_GCF_900444995.1
RS_GCF_900113665.1
Eremiobacteriales
Planctomycetota
Actinobateriota
UBA4705
Eudoremicrobiaceae
Chloroflexota
Actinobateriota
Planctomycetota
Patescibacteria
Chloroflexota
Cyanobacteria
Armatimonadota
Dormibacterota
UPB13
Chloroflexota
UBA4705
Eudoremicrobiaceae
B
genome
GB_GCA_002407045.1
GB_GCA_900313515.1
GB_GCA_002423485.1
GB_GCA_002329445.1
OLPPLKCL
OCMKBGHM
LGBFILLL
PIAMPJPB
HLLJDLBE
GB_GCA_003161555.1
GB_GCA_003140215.1
GB_GCA_003168375.1
GB_GCA_003164045.1
GB_GCA_003164495.1
GB_GCA_003136355.1
GB_GCA_003151115.1
GB_GCA_003140855.1
GB_GCA_003167055.1
GB_GCA_003155175.1
GB_GCA_003158175.1
GB_GCA_003166915.1
GB_GCA_003167155.1
GB_GCA_003140835.1
GB_GCA_003133745.1
GB_GCA_003243975.1
GB_GCA_003134035.1
GB_GCA_003158195.1
GB_GCA_003134695.1
GB_GCA_003136655.1
GB_GCA_003156715.1
GB_GCA_003136895.1
GB_GCA_003244105.1
GB_GCA_002410925.1
GB_GCA_003244185.1
RS_GCF_000023265.1
RS_GCF_000011905.1
RS_GCF_002973605.1
RS_GCF_000018865.1
OCMKBGHM
PIAMPJPB
HLLJDLBE
GB_GCA_000392435.1
RS_GCF_000007925.1
GB_GCA_000317655.1
RS_GCF_900444995.1
RS_GCF_000724625.1
RS_GCF_000284115.1
RS_GCF_900113665.1
UBA4705
Eudoremicrobiaceae
Eremiobacterales
Planctomycetota
Armatimonadota
Actinobateriota
Cyanobacteria
Patescibacteria
Eudoremicrobiaceae
Chloroflexota
Armatimonadota
Chloroflexota
Actinobateriota
Dormibacterota
UBP13

**<u>Figure S13: Phylogeny of the duplicated marker gene COG0124. Related to Figure 4.</u>**

(A) Investigating the evolutionary history of duplicated single-copy marker genes (here COG124), we found consistent duplication across *Ca*. Eudoremicrobeaceae and the parent order UBP9, thus ruling out the duplication as a signal of contamination in the binning process. The different evolutionary history of the second copy of COG124 (right-hand side of the tree), with closer relationship to Actinobacteria suggests that introgression events (including before the UBP9 and *Ca*. Eudoremicrobiaceae split) could be the origin of the increased genome size and biosynthetic potential observed in *Ca*. Eudoremicrobiaceae. (B) Similar patterns can be found in the second duplicated marker gene (COG522), although duplication was not detected across all *Ca*. Eudoremicrobeaceae spp. representatives.

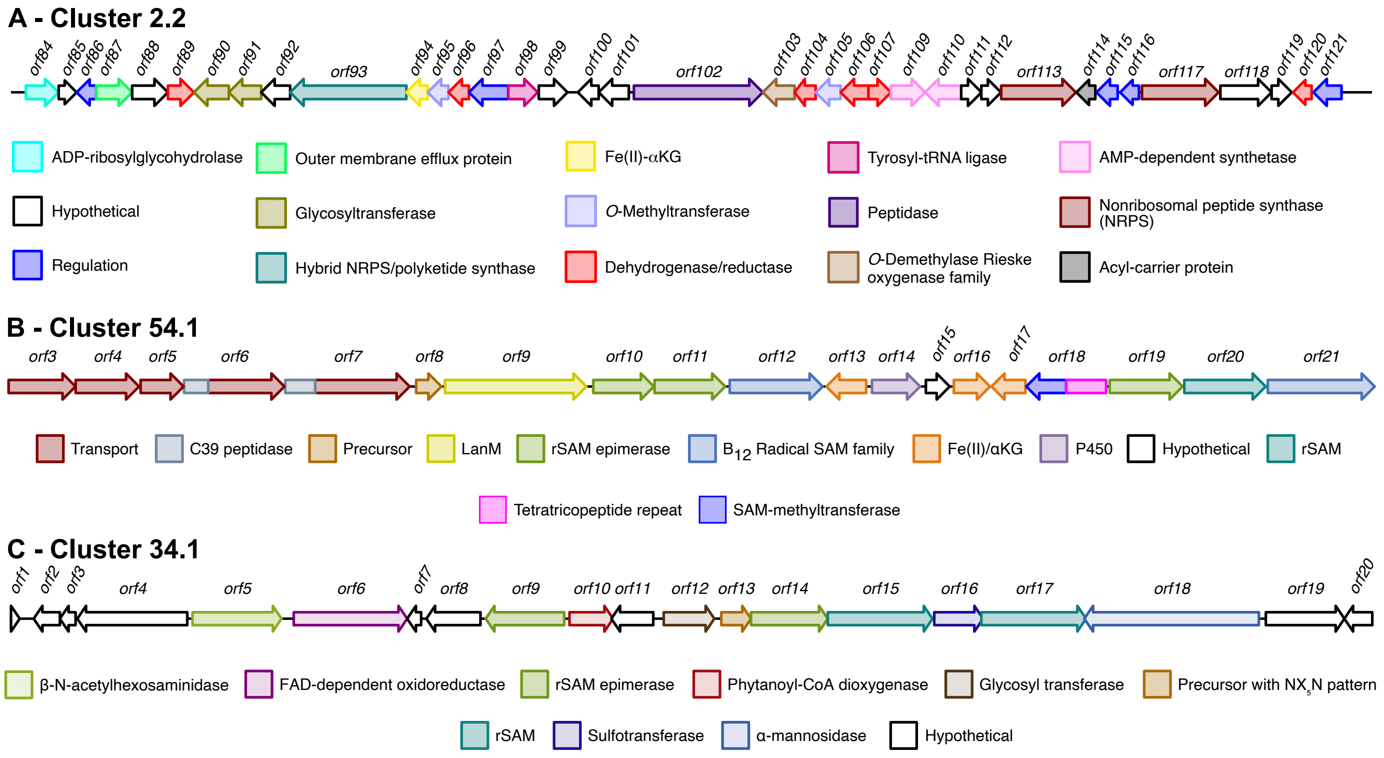
A - Cluster 2.2
orf84 orf85 orf86 orf87 orf88 orf89 orf90 orf91 orf92 orf93 orf94 orf95 orf96 orf97 orf98 orf99 orf100 orf101 orf102 orf103 orf104 orf105 orf106 orf107 orf109 orf110 orf111 orf112 orf113 orf114 orf115 orf116 orf117 orf118 orf119 orf120 orf121
ADP-ribosylglycohydrolase
Outer membrane efflux protein
Fe(II)-αKG
Tyrosyl-tRNA ligase
AMP-dependent synthetase
Hypothetical
Glycosyltransferase
O-Methyltransferase
Peptidase
Nonribosomal peptide synthase (NRPS)
Regulation
Hybrid NRPS/polyketide synthase
Dehydrogenase/reductase
O-Demethylase Rieske oxygenase family
Acyl-carrier protein
B - Cluster 54.1
orf3 orf4 orf5 orf6 orf7 orf8 orf9 orf10 orf11 orf12 orf13 orf14 orf15 orf16 orf17 orf18 orf19 orf20 orf21
Transport
C39 peptidase
Precursor
LanM
rSAM epimerase
B12 Radical SAM family
Fe(II)/αKG
P450
Hypothetical
rSAM
Tetratricopeptide repeat
SAM-methyltransferase
C - Cluster 34.1
orf1 orf2 orf3 orf4 orf5 orf6 orf7 orf8 orf9 orf10 orf11 orf12 orf13 orf14 orf15 orf16 orf17 orf18 orf19 orf20
β-N-acetylhexosaminidase
FAD-dependent oxidoreductase
rSAM epimerase
Phytanoyl-CoA dioxygenase
Glycosyl transferase
Precursor with NX5N pattern
rSAM
Sulfotransferase
α-mannosidase
Hypothetical

**Figure S14: Visual representations of BGCs encoded by *Ca*. E. malaspinii. Related to Figure 4.**

Visual representations and manual annotations of some *Ca*. Eudoremicorbium specific BGCs discussed in supplementary text, i.e. BGC 2.2 (A), BGC 54.1 (B) and BGC 34.1 (C). Color-coding corresponds to predicted enzyme domains and modifications. These can be interactively explored here:

https://microbiomics.io/ocean/db/1.0/marine_eremios/annotations/MALA_SAMN05422137_METAG_HLLJDLBE/antismash/MALA_SAMN05422137_METAG_HLLJDLBE-antismash/.

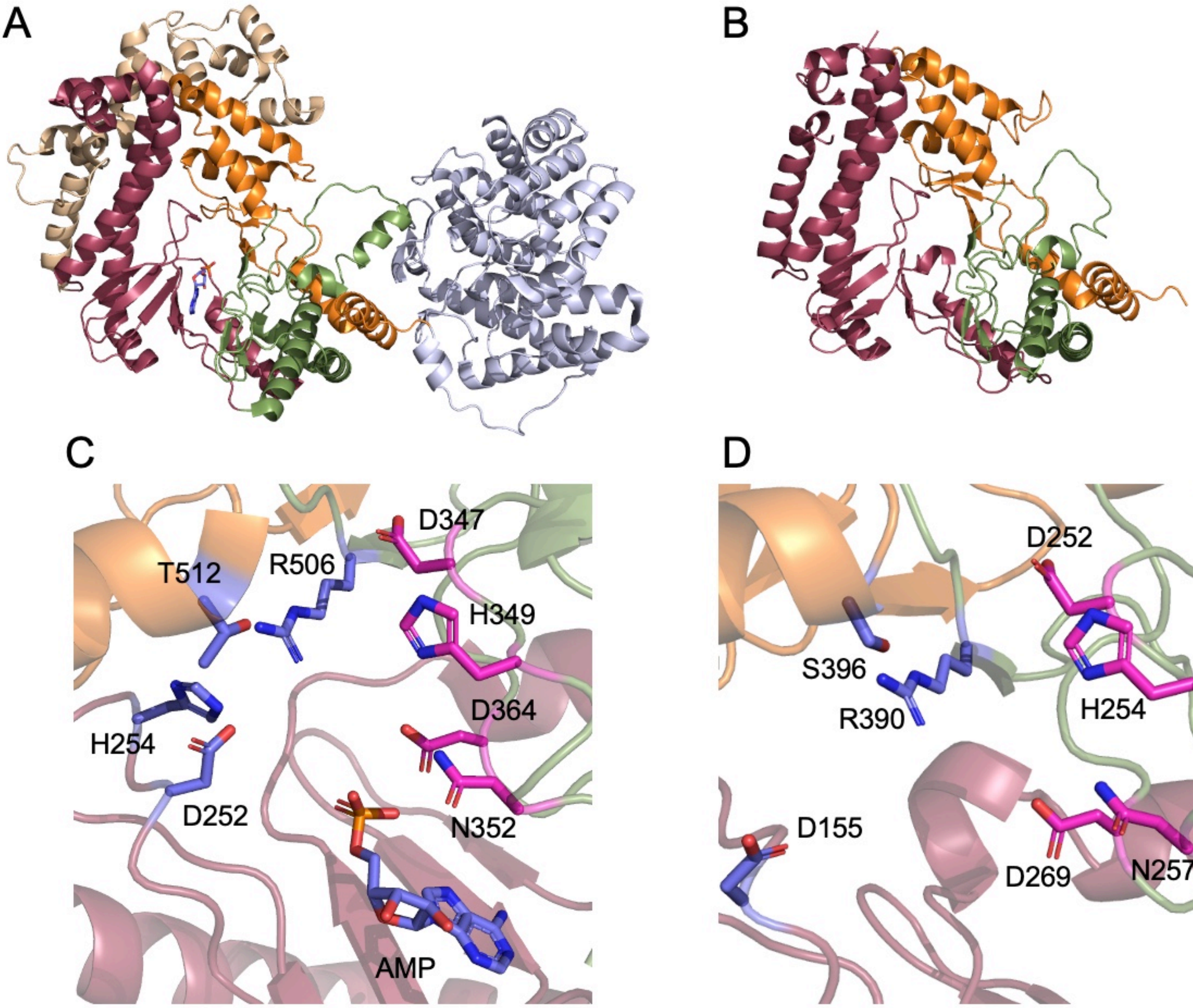

A
B
C
T512
R506
D347
H349
D364
H254
D252
N352
AMP
D
D252
S396
R390
H254
D155
D269
N257

**Figure S15: EmbM structural prediction and comparison to CylM (PDB: 5DZT). Related to Figure 4.**

(A) CylM crystal structure (Dong et al., 2015). Colored domains are involved in phosphorylation/dehydration and the domain in gray is responsible for cyclization. (B) EmbM structure prediction, highlighting similarities to CylM. (C) CylM active site. Residues in pink are proposed to be involved in phosphorylation and residues in purple are necessary for elimination. (D) Modelled active site of EmbM.

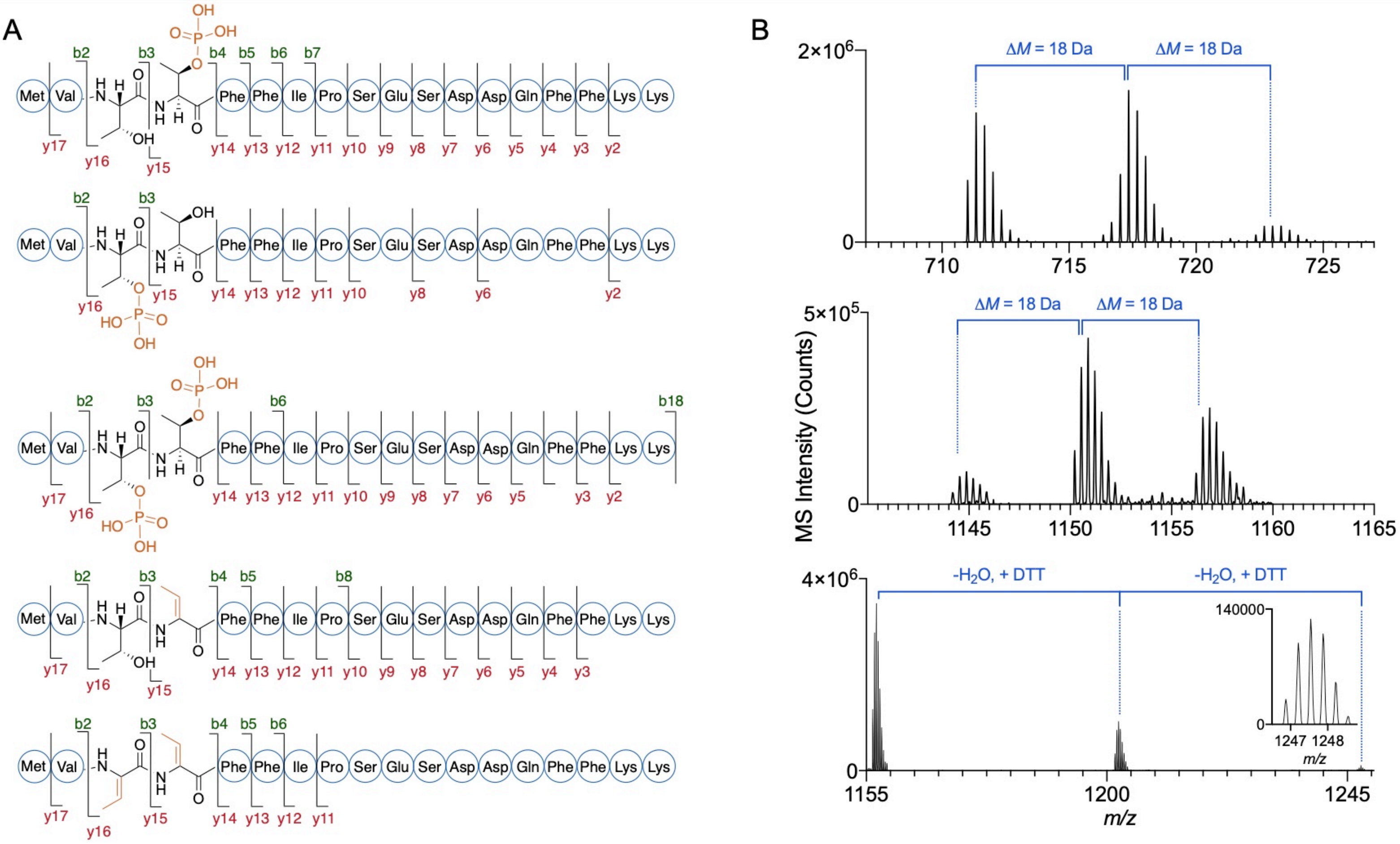

A
B
Met Val Phe Phe Ile Pro Ser Glu Ser Asp Asp Gln Phe Phe Lys Lys
b2 b3 b4 b5 b6 b7 b8 b18
y17 y16 y15 y14 y13 y12 y11 y10 y9 y8 y7 y6 y5 y4 y3 y2
ΔM = 18 Da
-H2O, + DTT
MS Intensity (Counts)
m/z
2×10^6
5×10^5
4×10^6
140000

**Figure S16: Mass spectrometry data for modified EmbA peptides. Related to Figure 4.**

(A) HR-MS/MS fragmentation of EmbA core at different modification stages (cleaved with LahT150). (B) Mass spectrum of dehydrated EmbA species: unmodified, single- and double dehydrated EmbA core (top); unmodified, single- and double dehydrated EmbA cleaved with trypsin (middle); and unmodified, single- and double dehydrated, DTT adduct of EmbA cleaved with trypsin (bottom).

### Supplementary Tables

**Table S1 - Sample accessions and metadata of the metagenomes and metatranscriptomes used in this study. Related to Figure 1.**

**Table S2 - Clustering the biosynthetic potential of the ocean microbiome. Related to Figure 2.**

**Table S3 - Discovery and description of the *Ca*. Eudoremicrobiaceae family. Related to Figure 4.**

**Table S4 - Transcriptomic analysis of natural *Ca*. E. taraoceanii populations. Related to FIgure 4.**

**Table S5 - Experimental characterization of a novel bacteriocin. Related to Figure 4.**
